# TM6SF2/PNPLA3/MBOAT7 loss-of-function genetic variants impact on NAFLD development and progression both in patients and in *in vitro* models

**DOI:** 10.1101/2020.12.13.422549

**Authors:** Miriam Longo, Marica Meroni, Veronica Erconi, Fabrizia Carli, Chiara Macchi, Francesco Fortunato, Dario Ronchi, Silvia Sabatini, Erika Paolini, Emilia Rita De Caro, Anna Alisi, Luca Miele, Giorgio Soardo, Giacomo Pietro Comi, Luca Valenti, Massimiliano Ruscica, Anna L Fracanzani, Amalia Gastaldelli, Paola Dongiovanni

**Affiliations:** General Medicine and Metabolic Diseases; Fondazione IRCCS Cà Granda Ospedale Maggiore Policlinico, Milan, Italy; Department of Clinical Sciences and Community Health, Università degli Studi di Milano, Milano, Italy; Department of Pathophysiology and Transplantation, Università degli Studi di Milano, Fondazione IRCCS Cà Granda Ospedale Maggiore Policlinico, Milan, Italy; Institute of Clinical Physiology, National Research Council (CNR), Pisa, Italy; Department of Pharmacological and Biomolecular Sciences, Università degli Studi di Milano, 20133 Milano, Italy; Research Unit of Molecular Genetics of Complex Phenotypes, “Bambino Gesù” Children’s Hospital IRCCS, Rome, Italy; Area Medicina Interna, Gastroenterologia e Oncologia Medica, Fondazione Policlinico Universitario A. Gemelli IRCCS, Rome, Italy; Department of Medical Area (DAME), University of Udine and Italian Liver Foundation, Bldg Q AREA Science Park - Basovizza Campus, Trieste, Italy; Neuromuscular and Rare Diseases Unit, IRCCS Foundation Ca’ Granda Ospedale Maggiore Policlinico, Milan, Italy; Trasfusional Center-Translational Medicine, Fondazione IRCCS Cà Granda Ospedale Maggiore Policlinico, Milan, Italy

**Author notes:** These authors equally contributed to the manuscript (Miriam Longo, Marica Meroni). **Author Contributions:** M.L. designed, wrote and revised the manuscript, collected, analyzed and interpreted data and prepared figures/tables; M.M. took part in the experimental design, data interpretation and reviewed the manuscript; V.E and E.P. contributed to *in vitro* study and reviewed the manuscript; P.P. provided TEM platform and supported data analysis; F.C performed lipidomic analysis and supported data analysis; S.S and ER.D.C. participated to bioinformatic analysis of lipidomic data; C.M, F.F., D.R. and GP.C. assessed mitochondrial functionality, supported data analysis and interpretation and reviewed the manuscript; L.M., G.S. L.V. were involved in data and samples collection; A.A., M.R., AL.F. and A.G. contributed to discussion, manuscript revision, data interpretation; P.D. took part to study design, manuscript drafting, data analysis and interpretation, study funding, supervision and has primary responsibility for final content. All the authors read and approved the final draft. **Competing Interests:** The authors declare no competing interests.

**Keywords:** NAFLD, HCC, TM6SF2, ER stress, mitochondrial dynamics

## Abstract

**Background and aims:** The I148M PNPLA3, the rs641738 in *MBOAT7/TMC4* locus and the E167K TM6SF2 polymorphisms represent the main predisposing factors to non-alcoholic fatty liver disease (NAFLD) development and progression. We previously generated a full knockout of *MBOAT7* in HepG2 cells (MBOAT7^-/-^), homozygous for the I148M PNPLA3. Therefore, we aimed to:1) investigate the synergic impact of the 3 at-risk variants on liver injury and hepatocellular carcinoma (HCC) in a large cohort of NAFLD patients;2) create *in vitro* models of genetic NAFLD by silencing *TM6SF2* in both HepG2 and MBOAT7^-/-^ cells.

**Methods:** NAFLD patients (n=1380) of whom 121 had HCC were stratified with a semi-quantitative score ranging from 0 to 3 according to the number of *PNPLA3*, *TM6SF2* and *MBOAT7* at-risk variants. *TM6SF2* was silenced in HepG2 (TM6SF2^-/-^) and MBOAT7^-/-^ (MBOAT7^-/-^TM6SF2^-/-^) through CRISPR/Cas9.

**Results:** In NAFLD patients, the additive weight of these mutations was associated with liver disease severity and increased risk to develop HCC. In HepG2 cells, *TM6SF2* silencing altered lipid composition and induced the accumulation of micro-vesicular LDs, whereas the MBOAT7^-/-^TM6SF2^-/-^ cells showed a mixed micro/macro pattern of LDs. *TM6SF2* deletion strongly affected endoplasmic reticulum (ER) and mitochondria ultrastructures thus increasing ER/oxidative stress. Mitochondrial number raised in both TM6SF2^-/-^ and MBOAT7^-/-^TM6SF2^-/-^ models, suggesting an unbalancing in mitochondrial dynamics and the silencing of both *MBOAT7* and *TM6SF2* impaired mitochondrial activity with a shift towards anaerobic glycolysis. MBOAT7^-/-^TM6SF2^-/-^ cells also showed the highest proliferation rate.

**Conclusions:** The co-presence of the 3 at-risk variants impacts on NAFLD course, in both patients and experimental models affecting LDs accumulation, mitochondrial functionality and metabolic reprogramming towards HCC.

## Introduction

Non-alcoholic fatty liver disease (NAFLD) is a growing burden on global healthcare, and it is considered the most relevant liver disease of the twenty-first century affecting both adults and children. It is predicted to become the leading cause of hepatocellular carcinoma (HCC) and the most common indication for liver transplantation by 2030^1^. NAFLD encompasses a wide spectrum of hepatic conditions ranging from simple steatosis (hepatic fat>5%) to nonalcoholic steatohepatitis (NASH), fibrosis, cirrhosis and HCC^2^. The pathogenesis of NAFLD is closely intertwined with increased adiposity, insulin resistance and dyslipidemia^3^. Besides environmental factors, the 50-70% of hereditable traits contributed to NAFLD susceptibility and its inter-individual phenotypic variability^4^. Three main single nucleotide polymorphisms (SNPs) have been identified in the *Patatin-like Phospholipase Domain-containing 3* (*PNPLA3),* the *Membrane Bound O-acyltransferase Domain-containing 7* (*MBOAT7*) and in the *Transmembrane 6 Superfamily Member 2* (*TM6SF2*) genes^5–7^ and have been associated with the NAFLD spectrum.

Intracellular fat accumulation and aberrant lipid metabolism represent the earliest events occurring in NAFLD and genetics may participate to hasten steatosis development and its transition to NASH and eventually to HCC^8^. PNPLA3 localizes on the surface of lipid droplets (LDs) and functions as triacylglycerols (TAGs) lipase. The rs738409 C>G variant in PNPLA3 encoding for 148 Isoleucine to Methionine aminoacidic substitution (I148M) inhibits TAGs hydrolysis and led to the accumulation of mutant PNPLA3 on LDs surface. MBOAT7 enzymatically remodels acyl-chains of phospholipids on cellular membranes by transferring polyunsaturated fatty acids to lyso-phosphatidylinositols (lyso-PI). The rs641738 C>T variant has been associated with the reduced hepatic expression of MBOAT7, determining changes in PI species towards saturated ones, precursors of TAGs synthesis and favoring fat accumulation^9^. Finally, TM6SF2 localizes in the endoplasmic reticulum (ER) and ER-Golgi compartments^10^ and it participates to TAG-rich lipoproteins’ lipidation and assembly in the ER cisternae^5, 11^. The rs58542926 C>T variant in the TM6SF2 gene encoding Lysine instead of Glutamate at residue 167 (E167K) causes the retention of very low-density lipoprotein (VLDL) in the liver and increases the intrahepatic TAGs content but it protects against cardiovascular complications^5, 12^. However, differently from I148M PNPLA3 and rs641738 *MBOAT7* variants, its role in cell injury and carcinogenesis remains uncharted.

It is well-established that the I148M PNPLA3, the rs641738 in *MBOAT7* and the E167K TM6SF2 SNPs predispose to NAFLD and advanced liver injury. Recently, it has emerged the opportunity to translate the genetics into clinics by aggregating these genetic variants in polygenic risk scores for the assessment of fatty liver development and progression^13^. However, the additive weight of the 3 at-risk mutations on liver disease severity and the related mechanisms need further investigations^14^. Therefore, we aimed to explore the synergic effects of the I148M PNPLA3, rs641738 *MBOAT7* and E167K TM6SF2 variants on clinical-pathological features and liver disease severity in a large cohort of patients with NAFLD. Moreover, to reproduce *in vitro* a condition which parallel human genetic NAFLD, we silenced in HepG2 hepatoma cells, homozygous for the I148M PNPLA3 variant, the *TM6SF2* and *MBOAT7* genes by exploiting CRISPR-Cas9 technology. We previously generated a full knockout of *MBOAT7* (MBOAT7^-/-^) in HepG2 cells which spontaneously developed LDs^9^. In this study, we silenced *TM6SF2* in both HepG2 (TM6SF2^-/-^) and MBOAT7^-/-^ (MBOAT7^-/-^TM6SF2^-/-^) cells in order to elucidate whether *TM6SF2* ablation, alone or in combination with that of *MBOAT7*, may induce pathological features resembling human NAFLD.

## Patients and methods

### Overall cohort

The Overall cohort consists of 1380 patients with NAFLD and it has been subdivided in the Hepatology Service cohort (n=1259) and the NAFLD-HCC cohort (n=121). Enrollment criteria, diagnosis and histological evaluation of the Overall cohort are described in the **Supplementary Materials.**

The Hepatology Service cohort and the NAFLD-HCC cohort were both stratified according to the number of variants as follows: 0 for patients who had no risk alleles; 1 for the presence of one risk allele heterozygous or homozygous in either *PNPLA3*, *MBOAT7* or *TM6SF2;* 2 for carriers who had two risk variants among *PNPLA3*, *MBOAT7* or *TM6SF2* in variable combinations;3 for subjects carrying all 3 at-risk variants either in heterozygous or homozygous. Demographic, anthropometric, and clinical features of the Overall cohort stratified according to class enrollment criteria or the number of PNPLA3 I148M, *MBOAT7* rs641738 and TM6SF2 E167K risk variants are shown in **Table S1-S2**.

### Gene silencing in HepG2 cells

Gene silencing of either *TM6SF2* or *MBOAT7* genes was induced by CRISPR-Cas9 in HepG2 hepatoma cells, carrying the I148M PNPLA3 mutation. We generated an inducible stable Cas9 cell line by lentiviral transfection in HepG2 cells (Cas9^+^). Then, *TM6SF2* was silenced in HepG2 and in *MBOAT7* knockout model (MBOAT7^-/-^)^9^ allowing us to obtain either a single (TM6SF2^-/-^) or a compound knockout stable model (MBOAT7^-/-^TM6SF2^-/-^) to study genetic NAFLD *in vitro*. Cas9^+^ cells wild type (Wt) in both *TM6SF2* and *MBOAT7* genes were used as control group (**Supplementary Materials**)

### Transmission Electron Microscopy (TEM)

LDs size was obtained by measuring at least 4 random diameters *per* LD through ImageJ software. The mean of diameters was used to calculate the average of both circumference (C=πd) and area (A=πR^2^) of LDs for each condition (**Table S3**). Quantification of ER width was obtained through ImageJ software by acquiring at least 3 measurements per ER lumen (**Table S4**). Mitochondria were counted by ImageJ software 15 random non-overlapping micrographs (bar scale:2 µm) containing at least a whole single cell (**Supplementary Materials**).

### Lipidomics

Lipid classes were separated with ultra-high pressure liquid chromatography (UHPLC) equipped with ZORBAX Eclipse Plus C18 2.1×100mm 1.8 µm columns and lipids concentrations were measured by MS-QTOF in both positive (for ceramides (Cers), phosphatidylcholines (PCs), Lyso-PCs, diacylglycerols (DAGs), TAGs and negative electrospray ionization (for PIs). Lipid species were quantified by Mass Hunter Profinder software based on the accurate mass m/z, as [M+1] ion, retention time and ion abundance and subtracting natural reference abundance (measured in the NIST quality standard) (**Supplementary Materials**).

### Statistical analysis

For descriptive statistics, continuous variables were reported as mean and standard deviation or median and interquartile range for highly skewed biological variables. Variables with skewed distribution were logarithmically transformed before analyses. The frequencies for each risk variants subgroup were compared using chi-squared (χ^2^) test. Differences between groups were calculated by one-way nonparametric ANOVA (Kruskal-Wallis), followed by post hoc *t-test* (two-tailed) when two groups were compared or Dunn’s multiple comparison test when multiple groups were compared adjusted for the number of comparisons. Lipidomic analyses were performed using the R package (**Supplementary Materials**). P values <0.05 were considered statistically significant. Statistical analyses were performed using JMP 14.0 (SAS, Cary, NC) and Prism software (version 6, GraphPad Software)

## Results

### The I148M PNPLA3, the rs641738 MBOAT7 and the E167K TM6SF2 genetic variants have a synergic effect on liver damage

In the Overall cohort, 172 patients were wild type (12.46%), 574 were heterozygous or homozygous for the I148M PNPLA3, the rs641738 *MBOAT7* or the E167K TM6SF2 (42.03%), 552 carried at least two different risk SNPs in variable combinations (40%) and 82 had all 3 at-risk variants (5.94%). In the NAFLD-HCC cohort the percentage of patients who carried 2 or 3 risk variants was higher compared to the Hepatology Service cohort (52% and 11.5% *vs* 38.84% and 5.4% respectively, p=0.004 and p=0.006; **Table S1**).

At generalized linear model adjusted for age, sex, BMI and type 2 diabetes (T2D), the co-presence of the 3 risk variants in the Overall cohort was associated with increased levels of markers of liver damage (p=0.003, β=0.07, 95% c.i.-0.02-0.11 and p<0.0001, β=0.07, 95% c.i.-0.04-0.11; **Table S2**). At ordinal logistic regression analysis adjusted as above the co-presence of the 3 at-risk variants associated with a higher grade of steatosis (p<0.0001, β=0.40, 95% c.i.0.27-0.55, **Figure 1A**), lobular inflammation (p<0.0001, β=0.29, 95% c.i.-0.15-0.44, **Figure 1B**), ballooning (p=0.004, β=0.25, 95% c.i. 0.07-0.42, **Figure 1C**), fibrosis (p<0.0001, β=0.42, 95% c.i.0.28-0.56, **Figure 1D**) and NAS score (p<0.0001, β=0.37 95% c.i.0.24-0.51, **Figure S1A**) in the Overall cohort. At nominal logistic regression analysis adjusted for age, sex, BMI and T2D, carriers of the 3 SNPs had an increased risk to develop NAFLD (OR: 1.40, 95% c.i. 1.06-1.83, p=0.01, **Figure S1B**), NASH (OR: 1.53, 95% c.i. 1.30-1.78, p<0.0001, **Figure 1E**), fibrosis >1 (OR: 1.57, 95% c.i. 1.30-1.89, p<0.0001, **Figure S1C**), fibrosis >2 (OR: 1.54, 95% c.i. 1.22-1.95, p=0.0003, **Figure S1D**), cirrhosis (OR: 1.62, 95% c.i. 1.22-2.14, p=0.0007, **Figure 1F**) and ~2-fold higher risk to develop HCC (OR: 1.73, 95% c.i.1.09-2.74, p=0.01, **Figure 1G**) even after the adjustment for the presence of fibrosis. Additionally, at bivariate analysis we found that the prevalence of the 3 risk variants was ~2.5 fold enriched in patients of the NAFLD-HCC cohort compared to those of the Hepatology Service cohort (p<0.0001, **Figure 1H**).

**Figure 1:**
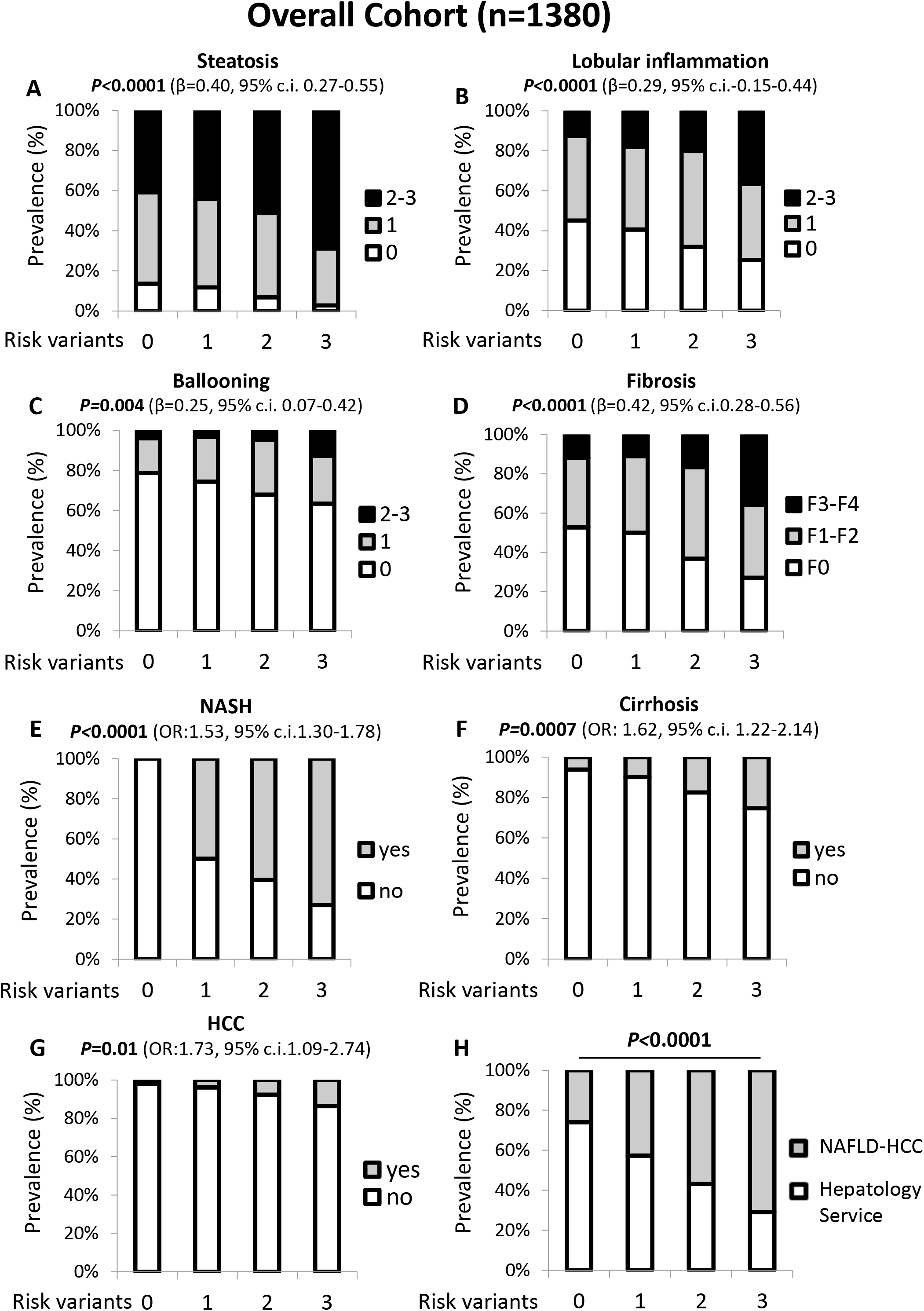
The co-presence of the PNPLA3 rs738409, MBOAT 7 rs641738 and TM6SF2 rs58542926 variants correlated with NAFLD spectrum. **A-D**) At ordinal regression analysis adjusted for age, gender, BMI and T2D, the co-presence of I148M PNPLA3, *MBOAT7* rs641738 and E167K TM6SF2 SNPs was associated with steatosis, lobular inflammation, ballooning and fibrosis stage. **E-G**) The increasing number of the at-risk variants correlated with NASH and cirrhosis at nominal logistic regression analysis adjusted for age, gender, BMI and T2D and with HCC after further adjustment for the presence of fibrosis. **H**) The co-presence of the 3 SNPs was significantly enriched the NAFLD-HCC cohort *vs* the Hepatology Service cohort (p<0.0001). *0: indicates the absence of risk variants; 1-2-3 indicates the total number of risk variants carried.

### TM6SF2 deletion alters lipid droplets size in hepatocytes

CRISPR/Cas9 system mediated *MBOAT7* and/or *TM6SF2* silencing further abrogating TM6SF2 function in lipoproteins export (**Figure S2A-E, Supplementary Materials**).

Then, we investigated whether genetically edited clones could reliably reproduce *in vitro* human steatosis. We assessed intracellular fat content through Oil Red O (ORO) staining. Consistently with our recent findings^9^, MBOAT7^-/-^ cells spontaneously accumulated giant LDs (**Figure 2A**, top; **Figure S3A-B**). Here, we found that TM6SF2^-/-^ cells developed small LDs at baseline, whereas the MBOAT7^-/-^TM6SF2^-/-^ clones presented a mixed pattern with either large or small LDs (**Figure 2A**, top; **Figure S3A-B**), thus suggesting that *TM6SF2* loss-of-function may diversely affect LDs formation compared to that exerted by *MBOAT7* deletion. According to the qualitative results, measurement of ORO positive areas and intracellular TAGs revealed ~30-40 fold higher enrichment of lipids in TM6SF2^-/-^ and MBOAT7^-/-^TM6SF2^-/-^ cells compared to control (p=0.0002 at ANOVA, adjusted p<0.01 *vs* Cas9^+^, **Figure 2B-C**), while in the MBOAT7^-/-^ ones the increase in fold was ~50, supporting that *MBOAT7* deletion exerted the largest influence on lipid handling.

**Figure 2:**
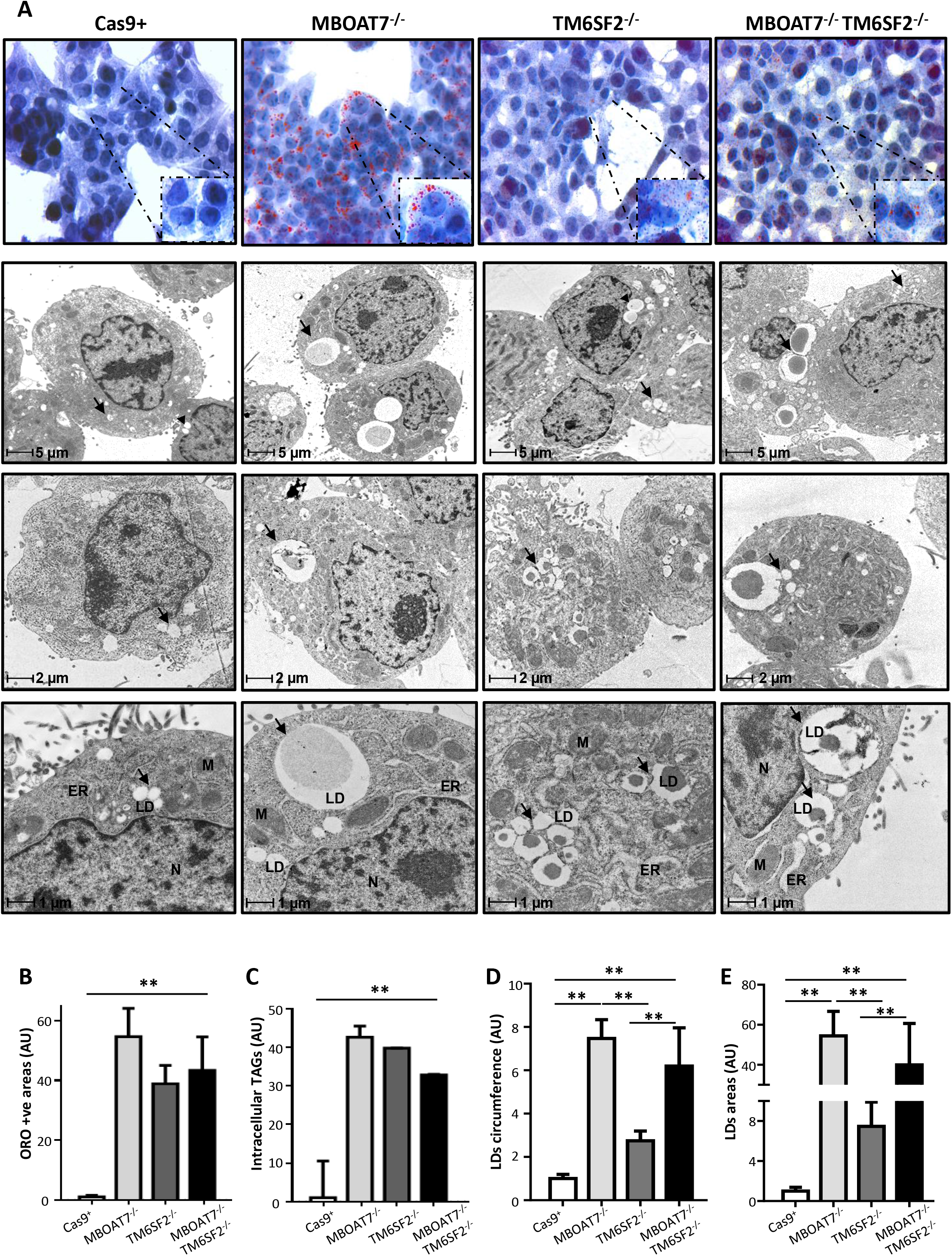
TM6SF2 deficiency induces small LDs budding in hepatocytes. **A**) On top, spontaneous development of LDs in TM6SF2^-/-^ and MBOAT7^-/-^ TM6SF2^-/-^ cells assessed by ORO staining (630X magnification). On bottom, representative TEM images of LDs obtained by ultrathin 70 nm sections of hepatocytes (bar scale: 1-2-5 μm). Black arrows indicated LDs. Capital letters indicates nucleus (N), endoplasmic reticulum (ER), mitochondria (M). **B**) ORO positive (+ve) areas were quantified by ImageJ in 10 random non-overlapping micrographs *per* condition by calculating the percentage of pixels above the threshold value in respect of total pixels *per* area. **C**) Measurement of TAGs content in cell lysates (adjusted **p<0.01 *vs* Cas9^+^). **D-E**) The average of LDs’ circumference and area was calculated from 10 random non-overlapping micrographs (bar scale: 1 μm) by ImageJ. Data are expressed as fold increase (Arbitrary Unit-AU) *vs* control. At least three independent experiments were conducted.

Therefore, we deeply examined intracellular fat content through high-resolution TEM imaging, which highlighted remarkable differences in LDs size (**Figure 2A**, bottom; **Figure S3C**). Whilst Cas9^+^ cells showed scarce and quite small lipid bodies, undetectable with ORO staining, the MBOAT7^-/-^, TM6SF2^-/-^ and MBOAT7^-/-^ TM6SF2^-/-^ clones differed in fat deposits’ volumes. Indeed, MBOAT7^-/-^ showed the largest LDs circumference and area (median size: 6.51 µm^2^ *vs* 0.11 µm^2^, **Table S3;** adjusted p<0.01 *vs* Cas9^+^ and TM6SF2^-/-^, **Figure 2D-E**). Conversely, TM6SF2^-/-^ cells had clustered areas enriched in much smaller LDs (median size: 0.87 μm^2^ *vs* 0.11 μm^2^, **Table S3**; p<0.0001 at ANOVA, adjusted p<0.01 *vs* Cas9^+^ and MBOAT7^-/-^, **Figure 2A**, bottom; **2D-E**). MBOAT7^-/-^TM6SF2^-/-^ cells exhibit features in-between MBOAT7^-/-^ and TM6SF2^-/-^ cells showing areas with a mixed pattern of greater or smaller LDs (median size: 4.60 μm^2^ *vs* 0.11 μm^2^, **Table S3**; p<0.0001 at ANOVA, adjusted p<0.01 *vs* Cas9^+^, **Figure 2A**, bottom**; 2D-E**). These results supported that *MBOAT7* and *TM6SF2* differently impacting on lipid handling could lead to a diverse distribution of micro and macro-vesicles.

### TM6SF2 ablation alone or combined to MBOAT7 impacts on lipid composition

To investigate whether differences in LDs dimension are correlated with changes in lipid species, we performed a lipidomic analysis of each experimental group. Levels of saturated/monounsaturated TAGs (**Figure S3D**; **Suppl. Materials 1**) were laden in MBOAT7^-/-^ cells possibly due to enhanced *de novo* lipogenesis^9^. TM6SF2^-/-^ cells were strongly enriched in highly saturated DAGs (**Figure 3A**) and TAGs (**Figure 3B-C**; **Suppl. Materials 1**) and less in unsaturated TAGs compared to the control group (**Figure 3B** and **3D**; **Suppl. Materials 1**).

**Figure 3:**
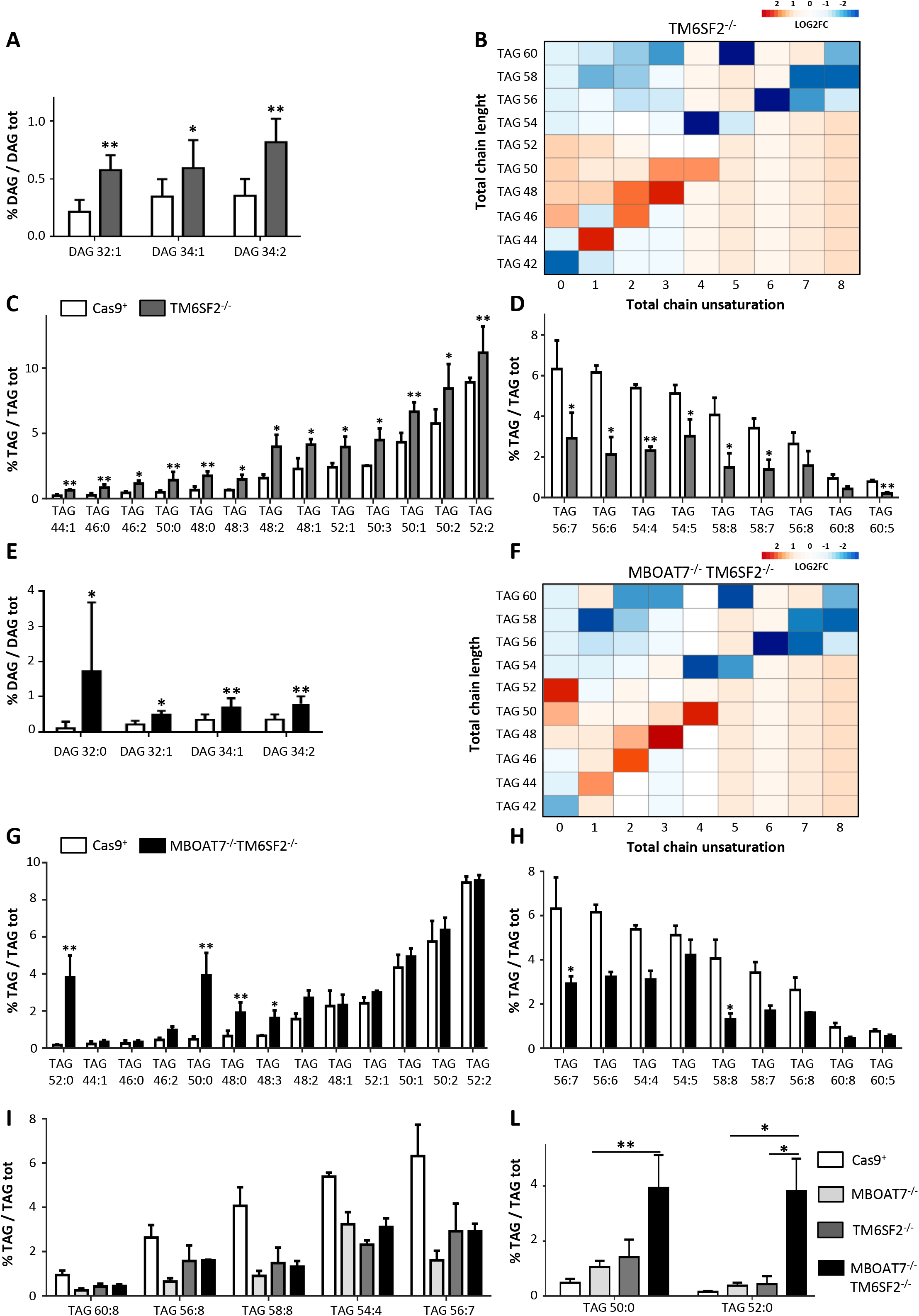
The impact of TM6SF2 deletion on LDs composition. **A-E**) Relatively enriched DAGs in TM6SF2^-/-^ and MBOAT7^-/-^ TM6SF2^-/-^ cells *vs* Cas9^+^. **B-F**) Heatmaps of TAGs were generated by calculating log2 fold change (log2FC) ratio between TM6SF2^-/-^/Cas9^+^ and MBOAT7^-/-^TM6SF2^-/-^/Cas9^+^ quantification. Red and blue boxes indicate over-expression or repression, respectively. **C-D-G-H**) Relatively enriched TAGs in TM6SF2^-/-^ and MBOAT7^-/-^ TM6SF2^-/-^ cells *vs* control. Data are expressed as percentage mean (%) of DAGs or TAGs and standard deviation (SD). adjusted *p<0.05 or p<0.01 *vs* Cas9^+^.

At principal component analysis (PCA), the MBOAT7^-/-^TM6SF2^-/-^ model showed an in-between lipid profile among MBOAT7^-/-^ and TM6SF2^-/-^ cells (**Figure S3E**). Similar to the single knockout cells, the compound knockout increased saturated/monounsaturated DAGs (**Figure 3E**) and TAGs (**Figure 3F-G**; **Suppl. Materials 1**) rather than long-chain polyunsaturated TAGs compared to control (**Figure 3H**; **Suppl. Materials 1**). Notably, we found that *TM6SF2* deletion rather than *MBOAT7* affected the amount of DAGs in the MBOAT7^-/-^TM6SF2^-/-^ cells by increasing mono/di-unsaturated DAG 32:1, DAG 34:1 and DAG 34:2 **(Figure S3F; Suppl. Materials 2**). Among the unsaturated TAGs, TAG 50:4, TAG 54:5, TAG 56:2 and TAG 58:7 were influenced by *TM6SF2* genetic background (**Figure S3G; Suppl. Materials 2**). However, the downregulation of most of the polyunsaturated TAGs did not show a prevailing impact between *MBOAT7* and *TM6SF2* deficiency **(Figure 3I**, **Suppl. Materials 1**).

In attempt to identify whether the MBOAT7^-/-^TM6SF2^-/-^ model may be characterized by peculiar TAG species, we found that the compound knockout markedly expressed the saturated TAG 50:0 and TAG 52:0 compared to Cas9^+^, MBOAT7^-/-^ and TM6SF2^-/-^ models (**Figure 3L Suppl. Materials 1; Suppl. Materials 2**), potentially mirroring the hepatic lipid profile of NAFLD patients and the most upregulated TAGs observed in HCC specimens.^15, 16^

### TM6SF2 silencing markedly induces ER stress in TM6SF2-silenced models

ER stress perturbs lipid metabolism, and it may play a crucial role in the NAFLD to NASH transition. Therefore, we assessed whether *TM6SF2* and *MBOAT7* deletion influenced ER morphology and function. We observed ultrastructural differences among each experimental group in the organization and width of ER lumen, signs of cellular stress. In normal condition, Cas9^+^ showed regular arrangement of ER cisternae, whose parallel tubules appeared continuous and randomly distributed throughout the cytoplasm (**Figure 4A**). In MBOAT7^-/-^ model, we found a significant enlargement of the ER lumen but still preserving its architecture (median width: 0.18 μm *vs* 0.09 μm, **Table S4**; p<0.0001 at ANOVA, adjusted p<0.01 *vs* Cas9^+^, **Figure 4A-B**) and increased GRP78 mRNA expression (p=0.0002 at ANOVA, adjusted p<0.05 *vs* Cas9^+^, **Figure 4C-D**), suggesting that a mild unfolded protein response (UPR) is associated with *MBOAT7* deletion as a possible consequence of intracellular fat content.

**Figure 4:**
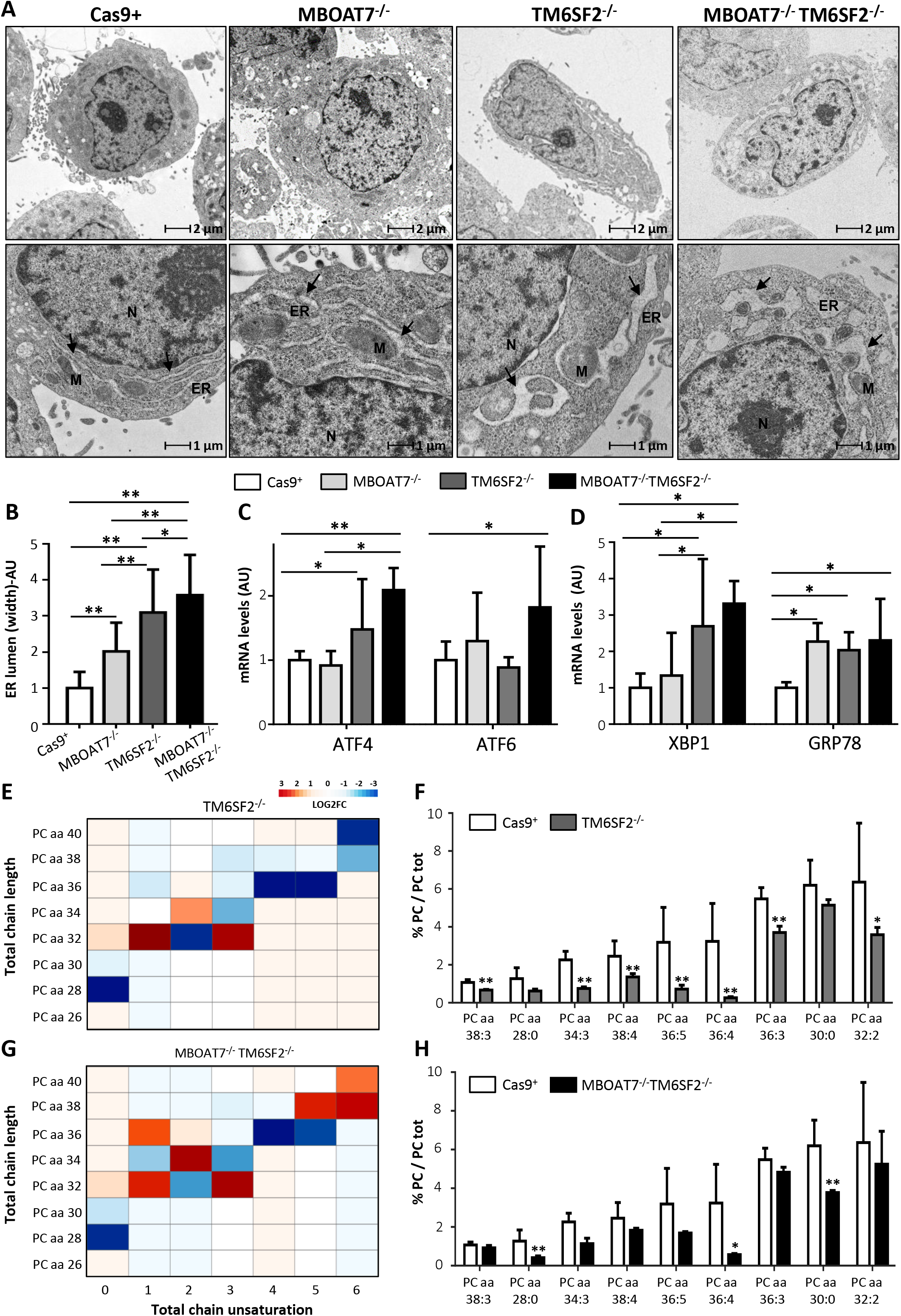
TM6SF2 silencing affects ER stress, morphology, and ER membranes’ fluidity. **A**) Representative TEM images of ER cisternae obtained by ultrathin 70 nm sections of hepatocytes (bar scale: 1-5 μm). Black arrows indicated ER tubules. Capital letters referred to nucleus (N), lipid droplets (LDs), mitochondria (M). **B**) ER width was obtained by taking at least 3 measurements *per* ER lumen (n=15 non-overlapping micrographs for each condition, bar scale: 1μm). **C-D**) The expression of ATF4/6, XBP1 GRP78 was evaluated by qRT-PCR and normalized to β-actin housekeeping gene. **E-G**) Heatmaps of PCs were generated by calculating log2 fold change (log2FC) ratio between TM6SF2^-/-^/Cas9^+^ and MBOAT7^-/-^ TM6SF2^-/-^/Cas9^+^ quantification. Red and blue boxes indicate overexpression or repression, respectively. **F-H**) Relatively enriched PCs in TM6SF2^-/-^ and MBOAT7^-/-^TM6SF2^-/-^ cells *vs* control. For qRT-PCR, data are expressed as fold increase (Arbitrary Unit-AU) *vs* control group. For lipidomic analysis, data are expressed as percentage mean (%) of PC species and standard deviation (SD). At least three independent experiments were conducted. adjusted *p<0.05 or **p<0.01 *vs* Cas9^+^ or *vs* MBOAT7^-/-^.

Notably, ER lumen appeared remarkably dilatated in cells lacking *TM6SF2* gene (median width: 0.30 and 0.35 μm *vs* 0.09 and 0.18 μm, **Table S4**), showing extremely disorganized tubules and fragmented cisternae compared to both Cas9^+^ and MBOAT7^-/-^ ones (p<0.0001 at ANOVA, adjusted p<0.01 *vs* Cas9^+^ and MBOAT7^-/-^, **Figure 4A-B**). According to these morphological changes, TM6SF2^-/-^ clones strongly upregulated markers of ER stress and UPR as ATF4 (p=0.0008 at ANOVA, p<0.05 *vs* Cas9^+^, **Figure 4C),** XBP1 (p=0.0001 at ANOVA, adjusted p<0.05 *vs* Cas9^+^ and MBOAT7^-/-^, **Figure 4D**) and GRP78 (p=0.0002 at ANOVA, adjusted p<0.05 *vs* Cas9^+^, **Figure 4D**). MBOAT7^-/-^TM6SF2^-/-^ cells showed an even more exacerbated breakage and enlargement of ER tubules, high local curvature of ER membranes (**Figure 4A-B**), and increased mRNA levels of ATF4 (p=0.0008 at ANOVA, adjusted p<0.05 *vs* MBOAT7^-/-^ and p<0.01 *vs* Cas9^+^, **Figure 4C**), ATF6 (p=0.03 at ANOVA, adjusted p<0.05 *vs* Cas9^+^, **Figure 4C**), XBP1 (p=0.0001 at ANOVA, adjusted p<0.05 *vs* Cas9^+^ and MBOAT7^-/-^, **Figure 4D**) and GRP78 (p=0.0002 at ANOVA, adjusted p<0.05 *vs* Cas9^+^, **Figure 4D**).

### Lacking TM6SF2 gene affects phosphatidylcholines’ metabolism leading to ER shapeless

Phospholipids exert a central role to maintain ER functions, membranes ‘fluidity and ER-mitochondria contact sites. Deletion in the *MBOAT7* gene reduced the amount of PC conjugated with arachidonoyl-CoA (*i.e.* PC 36:4, PC 38:6, PC 40:6, **Figure S4A**; **Suppl. Materials 1**) reinforcing previous data from our group^9^. Here, we found that *TM6SF2* silencing caused both Lyso-PCs and PCs depletion with either high or low side-chain saturation grade (**Figure 4E-H**; **Figure S4B-C**; **Suppl. Materials 1**) in both single and double knockouts compared to Cas9^+^. In particular, the compound knockout model dramatically reduced levels of saturated Lyso-PC 14:0, PC 28:0 and PC 30:0 compared to both control (**Figure S4B; Figure 4G-H**; **Suppl. Materials 1**) and MBOAT7^-/-^ (**Figure S4C-D, Suppl. Materials 2**) and this effect was amenable to *TM6SF2* shortage. Such evidence supports the pivotal role of *TM6SF2* deletion to drive ER morphological alterations and, in the double knockout, ER stress may be additively worsened by the presence of *MBOAT7* ablation.

### TM6SF2 silencing influences mitochondrial morphology, number and oxidative stress

Since mitochondrial dysfunction is a hallmark of human hepatic steatosis and its progression, we analyzed mitochondrial morphology and investigated oxidative injury in our *in vitro* models. At TEM, we observed features suggestive of mitochondrial damage and derangement in all knockout cell lines. In the control group, we found normo-shaped mitochondria with densely packed cristae, many of which longitudinally oriented while few others smaller in size probably resulting from mitochondrial fusion-fission balancing (mitobiogenesis) (**Figure 5A**). In MBOAT7^-/-^ cells, we found swollen and irregular mitochondrial cristae but still maintaining a quite normal morphology (**Figure 5A**). Conversely, both TM6SF2^-/-^ and MBOAT7^-/-^ TM6SF2^-/-^ cells displayed several areas enriched in mitochondria with small and globular shape, loss of cisterns’ architecture and ultrastructural electron density which may indicate mitochondrial failure and degeneration (**Figure 5A**).

**Figure 5:**
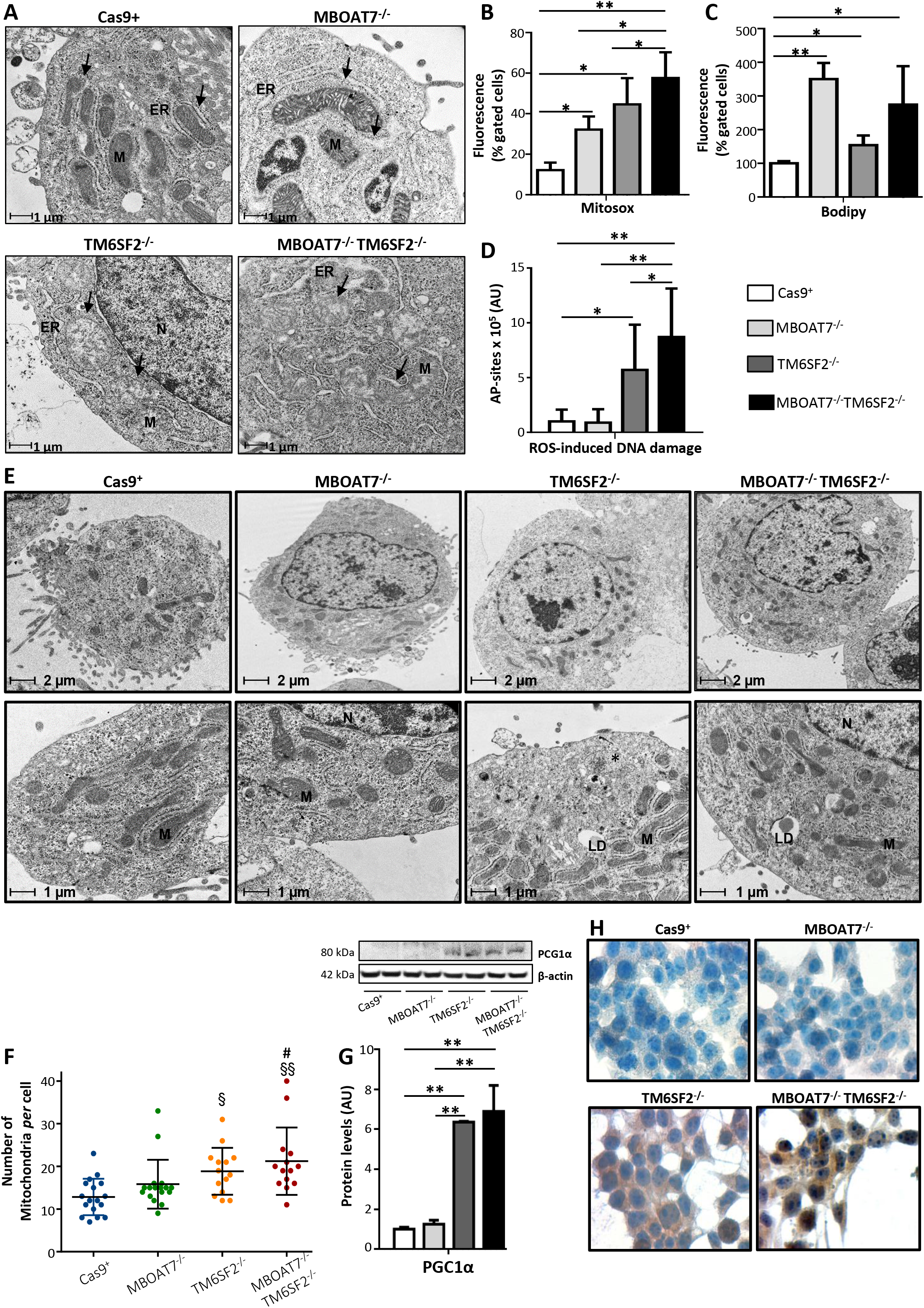
Alterations of mitochondrial degeneration and renewal in TM6SF2 knockout models. **A**) Representative TEM images of degenerative mitochondria (M) obtained by ultrathin 70 nm sections of hepatocytes (bar scale: 1 μm) are indicated by black arrows. **B-C**) ROS and lipid peroxides were measured in live cells through MitoSOX™ Red reagent and BODIPY 581/591 C11, respectively. **D**) Number of AP sites was obtained by isolating total DNA from each model. **E**) Representative TEM images of mitochondrial biomass obtained by ultrathin 70 nm sections of hepatocytes (bar scale: 5-1μm). Capital letters referred to nucleus (N), lipid droplets (LDs), endoplasmic reticulum (ER) and mitochondria (M). **F**) The number of mitochondria *per* cell was counted from 15 random non-overlapping micrographs (^§^p<0.05 and ^§§^p<0.01 *vs* Cas9^+^; ^#^p<0.05 *vs* MBOAT7^-/-^; bar scale: 2 μm). **G)**PGC1α protein levels were assessed by Western Blot and normalized to β-actin (**H**) Cytoplasmatic and nuclear localization of PGC1α protein was assessed in *in vitro* models. Adjusted *p<0.05 and **p<0.01 *vs* Cas9^+^ or *vs* MBOAT7^-/-^.

In keeping with these results, MBOAT7^-/-^, TM6SF2^-/-^ and MBOAT7^-/-^TM6SF2^-/-^ models hugely boosted ROS production compared to Cas9^+^ (p<0.0001 at ANOVA; adjusted p<0.05 *vs* control; **Figure 5B** and **Figure S5A**) and it was even worsened in the compound knockout (adjusted p<0.05 *vs* MBOAT7^-/-^ and TM6SF2^-/-^; **Figure 5B** and **Figure S5A**). The latter augmented the expression of MnSOD2 (p=0.002 at ANOVA, p<0.01 *vs* Cas9^+^ and MBOAT7^-/-^; **Figure S5B**) possibly as a compensatory mechanism to dampen ROS overflowing. We dug deeper into ROS-induced cellular damage and found that all mutated models exhibited a conspicuous raise in lipid peroxidation (p<0.0001 at ANOVA, adjusted p<0.05 *vs* Cas9^+^, **Figure 5C**; p=0.002 at ANOVA, p<0.05 *vs* Cas9^+^; **Figure S5C**). Furthermore, TM6SF2^-/-^ remarkably increased apurinic/apyrimidinic sites (AP), the main ROS-induced DNA damage (p<0.05 *vs* Cas9^+^, **Figure 5D**), whose levels were higher in the double knockout (p<0.01 *vs* Cas9^+^ and MBOAT7^-/-^, **Figure 5D**). Lower PPARα expression was observed only in the compound knockout (p<0.0001 at ANOVA; adjusted p<0.01 *vs* Cas9^+^, p<0.05 *vs* MBOAT7^-/-^ and TM6SF2^-/-^ cells, **Figure S5D**) supporting that the co-existence of *MBOAT7* and *TM6SF2* loss-of-functions may jointly affect lipid synthesis and catabolism contributing to progressive damage.

In parallel with the presence of degenerated mitochondria, the abundance of morphologically normal organelles was significantly increased in TM6SF2^-/-^ (p=0.003 at ANOVA, adjusted p<0.05 *vs* Cas9^+^; **Figure 5E-F**) and MBOAT7^-/-^TM6SF2^-/-^ cells (p=0.003 at ANOVA, adjusted p<0.05 *vs* MBOAT7^-/-^ and p<0.01 *vs* Cas9^+^, **Figure 5E-F**) as well as PGC1α protein levels, master regulator of mitobiogenesis (p<0.01 *vs* Cas9^+^ and MBOAT7^-/-^, **Figure 5G**). At immunocytochemistry, PGC1α markedly localized in the cytoplasm and in several nuclei of TM6SF2^-/-^ and MBOAT7^-/-^TM6SF2^-/-^ clones compared to Cas9^+^ and MBOAT7^-/-^ cells (**Figure 5H**), supporting its activation in response to unbalancing in mitochondrial fusion-fission events.

### MBOAT7^-/-^TM6SF2^-/-^model switches oxidative respiration towards anaerobic glycolysis

To explore mitochondrial functionality in NAFLD models, we measured the cytochrome c oxidase I (MT-COX1) levels, the main mitochondrially DNA (mtDNA)-encoded subunit of the complex IV, which were normalized on succinate dehydrogenase complex flavoprotein subunit A (SDHA) levels, a nuclear-encoded subunit of complex II. In MBOAT7^-/-^ clones, the MT-COX-I/SDHA ratio was reduced compared to control (**Figure 6A**). In both TM6SF2^-/-^ and MBOAT7^-/-^TM6SF2^-/-^ cells, the MT-COX-I/SDHA ratio was even lower (p<0.05 and p<0.01 *vs* Cas9^+^ and MBOAT7^-/-^, **Figure 6A**). The MT-COX-I and SDHA expression was independently evaluated by ELISA, which confirmed the significant reduction of MT-COX-I/SDHA ratio in either TM6SF2^-/-^ clones (p<0.0001 at ANOVA; p<0.05 and p<0.01 *vs* Cas9^+^ and MBOAT7^-/-^; **Figure 6B**) and the double knockout cells (p<0.0001 at ANOVA; p<0.01 *vs* Cas9^+^ and MBOAT7^-/-^; **Figure 6B**).

**Figure 6:**
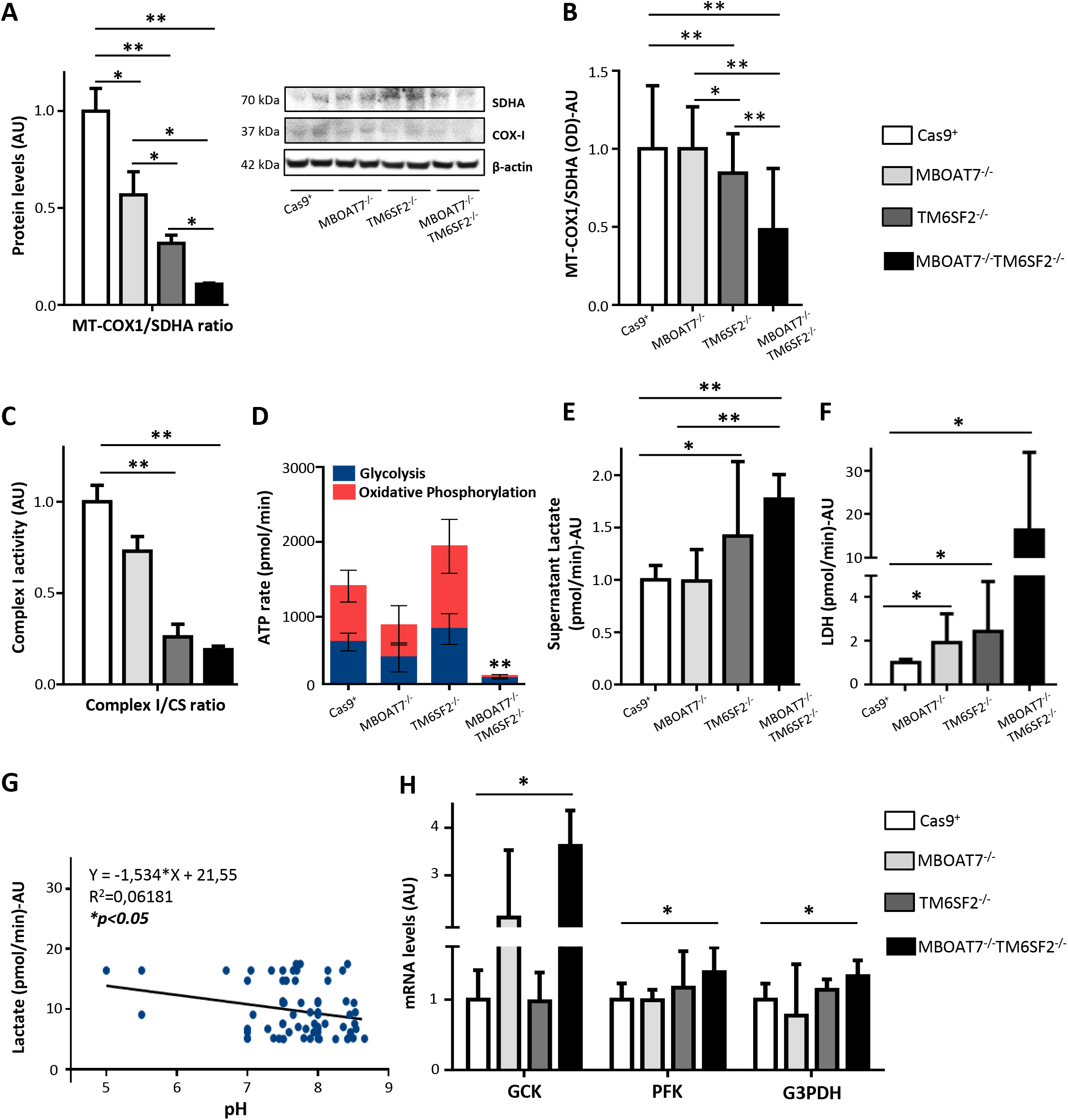
TM6SF2 silencing affects OXPHOS and when combined with MBOAT7 deletion, switches up to metabolic reprogramming. **A)** MT-COX1 levels were evaluated by Western Blot and normalized to SDHA **B)** MT-COX1 protein expression was measured by ELISA (λ=600 nm) and normalized to SDHA levels (λ=405 nm). **C**) Complex I enzymatic activity was biochemically evaluated and normalized to Citrate Synthase (CS). **D**) Total ATP rate was obtained by Seahorse XF Analyzers in live cells. **E-F**) Biochemical measurements of lactate (pmol/min) and LDH (pmol/min) were assessed in cell supernatants and lysates, respectively. **G**) Inverse correlation between secreted lactate levels and pH values. **H**) The mRNA levels of GCK, PFK, G3PDH were evaluated by qRT-PCR and normalized to β-actin housekeeping gene. Data are expressed as fold increase (Arbitrary Unit-AU) *vs* control. At least three independent experiments were conducted. adjusted *p<0.05 and **p<0.01 *vs* Cas9^+^ or *vs* MBOAT7^-/-^.

The enzymatic activity of complex I was significantly compromised in TM6SF2^-/-^ and MBOAT7^-/-^TM6SF2^-/-^ cells, suggesting that *TM6SF2* silencing may directly interfere with mitochondrial respiratory chain (p<0.05 and p<0.01 *vs* Cas9^+^ and MBOAT7^-/-^; **Figure 6C**). In keeping with these results, we provided a quantitative measurement of total ATP rate derived from mitochondrial and glycolytic pathways. We observed an equal contribution of mitochondria and glycolysis to total ATP production in all cell lines excluding the MBOAT7^-/-^TM6SF2^-/-^ ones. The latter significantly reduced the total amount of ATP (p<0.01 *vs* Cas9^+^, MBOAT7^-/-^ and TM6SF2^-/-^, **Figure 6D**), but further showed that the 87.1% of the ATP rate derived from glycolysis and only the 12.9% from oxidative phosphorylation (**Figure 6D**). Consistently, by measuring lactate production, the end-product of anaerobic glycolysis, in cell supernatants we found that TM6SF2^-/-^ clones increased lactate release and upregulated intracellular LDH which catalyzes the conversion of pyruvate to lactate and back (p<0.05 *vs* control, **Figure 6E-F**). No differences in lactate production were detected in the MBOAT7^-/-^ cells albeit it augmented LDH (p<0.05 *vs* control, **Figure 6E-F**).

In MBOAT7^-/-^TM6SF2^-/-^, lactate secretion was still more elevated than the other clones (p<0.01 *vs* Cas9^+^ and MBOAT7^-/-^, **Figure 6E**) and inversely correlated with pH values (**Figure 6G**). The compound knockout model further showed the highest intracellular LDH levels (~20-fold more than control; p<0.05 *vs* Cas9^+^; **Figure 6F**) and mRNA expression of glycolytic enzymes as GCK (adjusted p<0.05 *vs* Cas9^+^, **Figure 6H**), whose increasing levels were possibly due to the *MBOAT7* deficiency, PFK and G3PDH (adjusted p<0.05 *vs* Cas9^+^, **Figure 6H**), thus supporting the enhancement of glycolytic pathway.

In sum, we found that the combined silencing of *MBOAT7* and *TM6SF2* markedly impairs mitochondrial dynamics and run into metabolic reprogramming possibly contributing to progression to malignant transformation.

### MBOAT7^-/-^TM6SF2^-/-^ cells increase cell survival, proliferation, and invasiveness

In the hypothesis that *TM6SF2* and *MBOAT7* deletion in the background of *PNPLA3* mutation may sustain the carcinogenic phenotype, we evaluated growth potential and migration in our models. MBOAT7^-/-^, TM6SF2^-/-^ and the compound knockout hepatocytes had a higher proliferation rate compared to Cas9^+^ ones at 72 hours (adjusted p<0.01 *vs* control, **Figure 7A**). MBOAT7^-/-^TM6SF2^-/-^ clones further displayed the greatest growing ability upon 1 week (adjusted p<0.01 *vs* Cas9^+^, MBOAT7^-/-^ and TM6SF2^-/^, **Figure 7A**). At scratch assay, both MBOAT7^-/-^ and TM6SF2^-/-^ cells showed a greater wound healing capacity at 24 and 48 hours compared to Cas9^+^, whereas MBOAT7^-/-^ TM6SF2^-/-^ was able to almost completely repair the scratch after just 24 hours (**Figure 7B**) thereby showing the largest proliferative and invasiveness power. Consistently, TM6SF2-silenced cells aberrantly activated PI3K/Akt/mTOR cascade in absence of any *stimuli* (**Figure S6A-D**) and the compound knockout strongly delayed pharmacological response to Sorafenib, a multikinases inhibitor approved for the treatment of advanced HCC supporting its more proliferative and aggressive phenotype (**Figure S6E-F; Supplementary Materials**).

**Figure 7:**
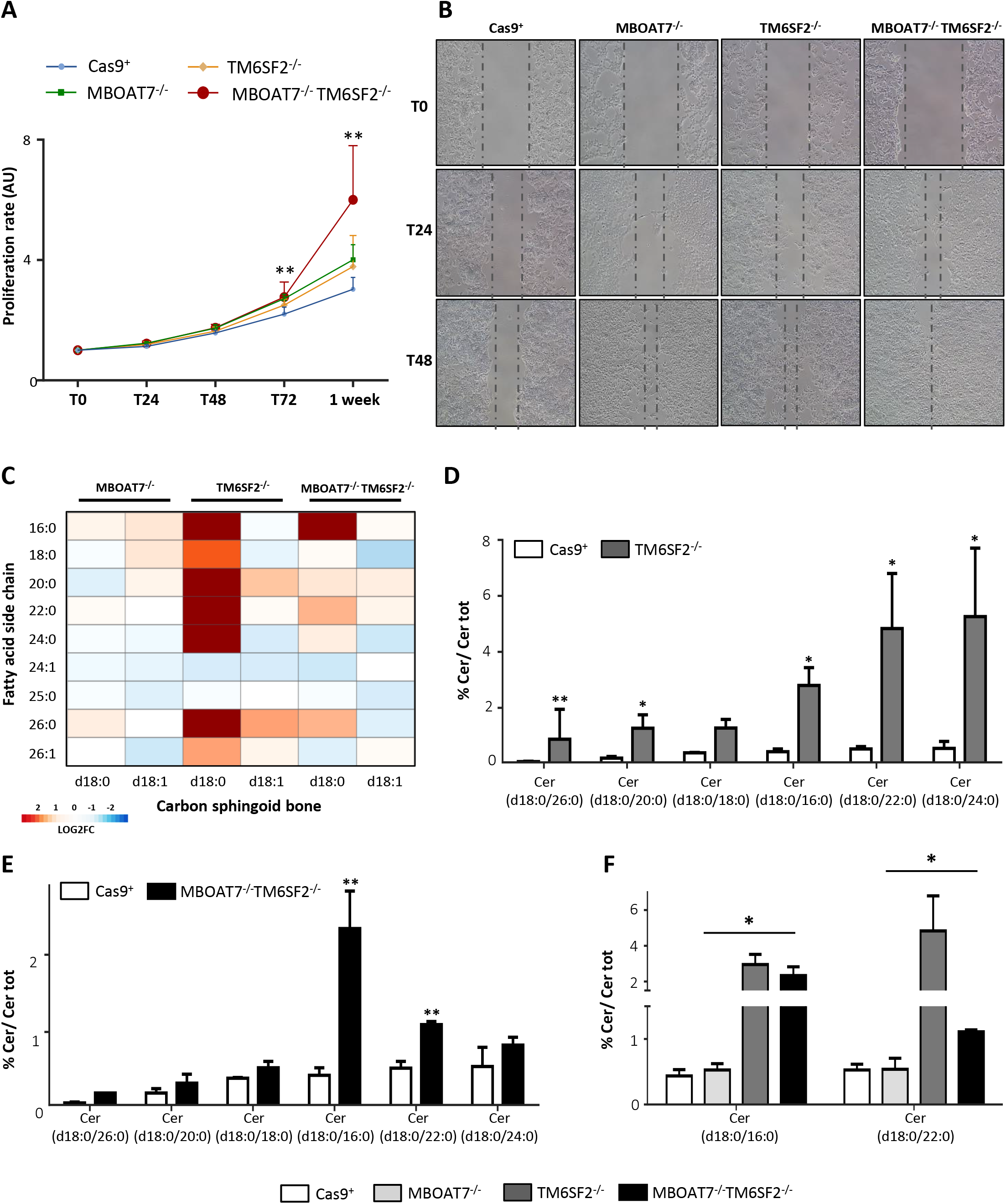
TM6SF2 lacking promotes cell survival and carcinogenesis. **A**) Cell growth was assessed through MTS assay for 24-48-72 hrs and 1 week (λ=490 nm). Data are expressed as fold increase *vs* control (Arbitrary Units-AU). **B**) Representative images of wound healing assay were acquired at 0-24-48 hrs (100X magnification). The dotted lines indicate the scratch width. **C**) Heatmap of dihydro-Cers was generated by calculating log2 fold change (log2FC) ratio between MBOAT7^-/-^/Cas9^+^, TM6SF2^-/-^/Cas9^+^ and MBOAT7^-/-^ TM6SF2^-/-^/Cas9^+^ quantification. Red and blue boxes indicate overexpression or repression, respectively. **D-E**) Relative abundance of dihydro-Cers in TM6SF2^-/-^ and MBOAT7^-/-^TM6SF2^-/-^ cells *vs* Cas9^+^. **F**) Relative abundance of dihydro-Cers in MBOAT7^-/-^TM6SF2^-/-^ cells *vs* MBOAT7^-/-^. Data are expressed as percentage mean (%) of Cer species and standard deviation (SD). adjusted *p<0.05 or **p<0.01 *vs* Cas9^+^ or *vs* MBOAT7^-/-^.

### Loss of TM6SF2 gene impacts on the abundance in dihydroceramide species

Dihydroceramides are precursors of ceramides and both exert different biological roles as modulators of cell fate. The percentage of dihydroceramides was strongly increased in the TM6SF2^-/-^ cells (**Figure 7C**), especially those species consisting of saturated sphingosine analog (d18:0) and incorporating long-chain fatty acids with no double bonds (adjusted p<0.05 and p<0.01 *vs* Cas9^+^; **Figure 7D**). An in-between sphingolipid phenotype was observed in the compound knockout model (**Figure 7C**), in which Cer (d18:0/16:0) and Cer (d18:0/22:0) showed the highest levels compared to Cas9^+^ (p<0.0001 at ANOVA, p=0.0002 and p=0.002, respectively; **Figure 7E**) and were probably driven by the *TM6SF2* rather the *MBOAT7* silencing (p<0.0001 at ANOVA, adjusted p=0.02 and p=0.01 *vs* MBOAT7^-/-^; **Figure 7F**). Therefore, the dramatic increase in dihydroceramides due to *TM6SF2* deletion is partially retained in the double knockout model and may contribute to the pro-survival phenotype observed in MBOAT7^-/-^TM6SF2^-/-^ cells.

## Discussion

In the present study, we carried out a comprehensive analysis in a large cohort of biopsy-proven NAFLD patients (n=1380), including 121 NAFLD-HCC individuals, in order to investigate the potential interactions among *PNPLA3*, *MBOAT7* and *TM6SF2* variants on liver disease severity and HCC development. Polygenic risk scores combining genetic predictors mainly associated to NAFLD have been proposed to improve diagnostic accuracy and to personalize therapeutic options^13^. Here, we showed that the co-presence of the I148M PNPLA3, rs641738 close to *MBOAT7* and E167K TM6SF2 SNPs significantly associated with liver function tests and histological degree of steatosis, lobular inflammation, hepatocellular ballooning, and fibrosis. Additionally, the risk to develop HCC in the Overall cohort was ~2 in presence of all 3 at-risk variants to the extent that the co-presence of the 3 SNPs was significantly enriched in NAFLD-HCC patients compared to those affected by NAFLD. Previously, the additive effect of E167K TM6SF2 and I148M PNPLA3 variants on hepatic steatosis was explored by Xu et al in 158 NAFLD subjects and matched controls, in whom fatty liver was non-invasively diagnosed by Fibroscan. They found that the prevalence of NAFLD was more than 5-fold higher in patients carrying both mutations compared to those with none or a single variant^17^. In a multicenter biopsy-based study including 515 patients with NAFLD who were genotyped for the 3 variants, Krawczyk et al showed a significant association between serum AST levels and the increasing number of risk alleles^18^ and the latter was further correlated to high HCC risk^14^.

We generated stable full knockouts of *MBOAT7*, *TM6SF2* or both genes in HepG2 cells, carrying the I148M PNPLA3 variant, through CRISPR-Cas9 in order to investigate the mechanisms related NAFLD pathogenesis in genetically-edited *in vitro* models. We previously demonstrated that *MBOAT7* deletion reduced MBOAT7 expression and affected its enzymatic activity leading to fat accumulation in HepG2 cells^9^ thus mirroring the condition observed in NAFLD carriers of the rs641738 T risk allele. Likewise, *TM6SF2* silencing alone or combined to the *MBOAT7* deficiency led to 50% reduction of TM6SF2 levels similarly to what reported by Ruhanen et al^19^, Smagris et al^11^ and O’Hare et al^20^, who induced stable *TM6SF2* knockdown in HuH7 hepatoma cells, hepatic *Tm6sf2* inactivation in mice and CRISPR/Cas9-mediated *tm6sf2* silencing in zebrafish, respectively. The role of TM6SF2 in lipoproteins’ metabolism, occurring in the ER-Golgi compartments, has been previously described in both experimental and clinical studies. In human hepatic 3D spheroids obtained from E167K donors, APOB-100 secretion was decreased^21^. Smagris and collaborators demonstrated that hepatic *Tm6sf2* deficiency impaired the “bulk phase” lipidation of VLDL^11^. According to these findings, our TM6SF2^-/-^ and MBOAT7^-/-^TM6SF2^-/-^ models suppressed APOB and TAG-rich lipoprotein release recapitulating features of patients carrying the E167K variant^12^.

To evaluate whether *TM6SF2* deletion alone or combined with the MBOAT7 one may reliably model *in vitro* human hepatic steatosis, we carried out an in-depth characterization of fat storing power and intracellular lipid profile. Both *MBOAT7* and *TM6SF2* silencing spontaneously developed LDs in hepatocytes further corroborating their involvement in steatosis onset. However, the different impact on LDs size according to the genetic background has never been reported in previous studies. *MBOAT7* deficiency induces giant LDs development, which resembles human macro-vesicular steatosis, by shunting saturated-PI towards synthesis of saturated/monounsaturated TAGs thus favoring DNL and perturbing membranes ‘dynamics^9^. Conversely, *TM6SF2* deletion may induce the formation of smaller, clustered LDs with a median size dimension of 0.87 μm^2^ assessed by TEM and that may remind micro-vesicular steatosis in humans. According to our data, Smagris and co-workers found that LDs distribution, evaluated through ORO staining, was detectable in the smallest size range consisting of 1-2 μm^2^ in liver sections of Tm6sf2^-/-^ mice^11^. Although further studies are required to investigate the role of micro-vesicular steatosis in the progression of liver disease, it has been demonstrated that the presence of micro rather than macro-steatosis correlated with hepatocellular ballooning, presence of Mallory-Denk bodies and mitochondrial dysfunction^22^. Notably, due to the contribution of both *MBOAT7* and *TM6SF2* loss-of-functions the compound knockout showed a mixed pattern of LDs content closely reflecting micro-macro vesicular steatosis which mainly characterizes liver histology of NAFLD patients. The lipidomic analysis revealed that TM6SF2-silenced cells mostly over-expressed TAGs incorporating saturated/monounsaturated fatty acid chains at the expense of polyunsaturated TAGs, reflecting the lipidomic data obtained in experimental models^19, 23^ and liver biopsies of E167K carriers^23^. Besides, the upregulation of TAG 50:0 and TAG 52:0, lipid biomarkers featuring serum and hepatic signature of NAFLD patients^15^ and, even more, in those arising HCC, were observed in the double knockout, thus supporting the deleterious effects synergically caused by the co-presence of *MBOAT7* and *TM6SF2* genetic deficiencies. Therefore, our findings supported by conspicuous evidence *in vitro*, *in vivo* and in humans suggested that TM6SF2^-/-^ cells may trustworthily be exploited as genetic NAFLD *in vitro* model as it fairly summarizes phenotypic traits of patients carrying the E167K variant. *TM6SF2* deletion induced the accumulation of micro-LDs compared to those developed by *MBOAT7* knockout cells and the mechanism underlying intracellular fat storage probably takes account of the increase in DAGs/TAGs with high degree of saturation and, overall, from reduction of polyunsaturated TAGs besides the retention of TAG-rich lipoproteins. As regards the MBOAT7^-/-^TM6SF2^-/-^ model, it showed a pattern of LDs volumes and lipid signature in-between the MBOAT7^-/-^ and TM6SF2^-/-^ thus representing the first model generated *in vitro* which may fully reproduce features of NAFLD individuals bearing all the 3 at-risk variants.

ER and mitochondrial dysfunction are hallmark of NAFLD as they may promote its progression to NASH and HCC. Genetic risk variants may actively participate to disease progression within the hepatic cells, although it has not been investigated whether and to which extent they can lead to organelles’ impairment. In the present study, we showed that ER tubules and mitochondrial cristae were modestly enlarged in the MBOAT7^-/-^ cells but still maintaining their morphological architecture and this condition was coupled to both ER/oxidative damage, probably resulting from lipotoxicity induced by the presence of large LDs.

The impact of *TM6SF2* silencing on ER morphology was only described by O’Hare et al^20^ in human enterocytes, in small intestine and liver of zebrafish larvae, in which TM6SF2 was acutely downregulated. Here, we found that the distance among ER tubules was more than 3-fold larger in cells lacking *TM6SF2* gene, which also presented enhanced ER stress and a dramatic reduction in PCs abundance. PC represent one of the main components of cellular membranes and their depletion has been associated with changes in ER morphology^24^ and, most recently, as hepatic lipid signature of NAFLD carriers the *TM6SF2* T risk allele^23^.

Consistently with the biological role of TM6SF2, inhibition of PC production affects the amount VLDL particles trafficking in the ER-Golgi compartments^25^ and it could be speculated that micro-LDs developed by TM6SF2^-/-^ and MBOAT7^-/-^TM6SF2^-/-^ models may arise from low PC abundance whose levels impact on LDs expansion^26^.

Errors in mitochondrial dynamics have been pointed out in NAFLD humans and experimental models as they may drive towards NASH and finally to metabolic reprogramming and malignant transformation leading to HCC. Notwithstanding, studies are still controversial, and none reported the effect of *MBOAT7* or *TM6SF2* silencing on mitobiogenesis. As aforementioned, MBOAT7^-/-^ cells affected mitochondrial morphology and oxidative stress as a possible result of the huge amount of intracellular fat accumulation but without altering number of mitochondria. Intriguing findings emerged in the *TM6SF2* knockout cells where both mitochondrial degeneration and high mitochondrial biomass came out, suggestive of an unbalance in fusion, fission and mitophagic processes. Alterations in mitochondrial lifecycle were also supported by increasing protein levels of PCG1α and may also take account to the loss of ER ultrastructure, which actively participate to mitochondrial dynamics, and OXPHOS activity thereby suggesting that *TM6SF2* downregulation may directly intervene to prompt mitochondrial dysfunction.

Changes in lipid composition (DAGs, TAGs, cholesterol esters) and rise in lactate production may be a sign of the *Warburg effect*, the metabolic shifting towards anaerobic glycolysis and mitochondrial derangement^27–29^. Min et al^30^ demonstrated that the overexpression of mutant I148M PNPLA3 in HuH-7 cells was associated to enhanced levels of lactate. Similarly, our study revealed that the double knockout showed additional aspects related to mitochondrial failure as it suppresses both β-oxidation and ATP production, whose levels arises for the 87.7% from anaerobic glycolysis, and showed the highest expression of glycolytic enzymes, lactate release and growth potential. Additionally, increased levels of dihydroceramides in the liver of obese NASH individuals have been correlated with mitochondrial failure^31^ and the ratio of dihydroceramides to ceramides appears crucial to determine cell fate^32, 33^. Only recently, Banini et al^34^ found that high levels of DAG, TAG and Cer species were associated to metabolic reprogramming in NAFLD rodents expressing the I148M mutation. Here, we firstly reported that *TM6SF2* silencing heavily enhanced dihydroceramides (d18:0) binding saturated long-chain fatty acids (*i.e* palmitic acid (16:0)), which is further consistent with the increase in shorter chain saturated DAGs and TAGs, and suggestive of a pro-survival phenotype. However, the co-presence of *MBOAT7* and *TM6SF2* deletion did not induce an even more increment of dihydroceramide content suggesting that *MBOAT7* may diversely contribute to the carcinogenesis in the double knockout. As occurs in carriers of all 3 at-risk alleles which exhibit the most elevated HCC prevalence, loss of both *MBOAT7* and *TM6SF2* synergistically promote an aggressive phenotype involving different mechanisms, but whether the I148M PNPLA3 genetic background of the HepG2 cells participate to metabolic switching cannot be ruled out.

To conclude, this study highlighted how *MBOAT7* and *TM6SF2* silencing diversely impact of lipid metabolism, ER/mitochondrial dynamics and tumorigenesis. These results may explain on one hand the impact of the I148M PNPLA3, *MBOAT7* rs641738 and E167K TM6SF2 variants on steatosis development and, on the other how they can promote the switch to NASH up to HCC. Moreover, we firstly proposed that *MBOAT7* and *TM6SF2* deletion may be decisive in developing a different macro and/or micro LDs pattern in hepatocytes. If *MBOAT7* deficiency predominantly induces large LDs formation by affecting PI metabolism with a mechanism boosting DNL, the small LDs observed in *TM6SF2*-silenced cells probably involve ER dysfunction and PCs depletion, which potentially affect LDs swelling. We also revealed that *TM6SF2* loss-of-function unbalances hepatic mitochondrial biogenesis, interferes with the activity of multi-enzymatic complexes of the respiratory chain thus extolling the oxidative and inflammatory status and accumulates dihydro-Cer species possibly enhancing its growth potential. In sum, we showed that the accrual of at-risk variants in *PNPLA3*, *MBOAT7* and *TM6SF2* genes predisposes to NAFLD and its progression towards cancer. Moreover, by exploiting a novel *in vitro* model we proposed some mechanisms through which these mutations impact on disease onset and severity. Notably, compound knockout showed matched characteristics of both single knockouts as regards lipid composition and organelles’ derangement, which together may contribute to the *Warburg effect* attempting to assume a pro-survival phenotype as occurs during hepatocarcinogenesis.

## Supporting information

Lipidomic data 1

Lipidomic data 2

## Acknowledgements

The authors would sincerely thank Dr. Podini Paola, IRCCS Ospedale San Raffaele, (Milan, Italy) for the technical support provided for the Transmission Electron Microscopy (TEM) and the intellectual contribution for data analysis interpretation.

---

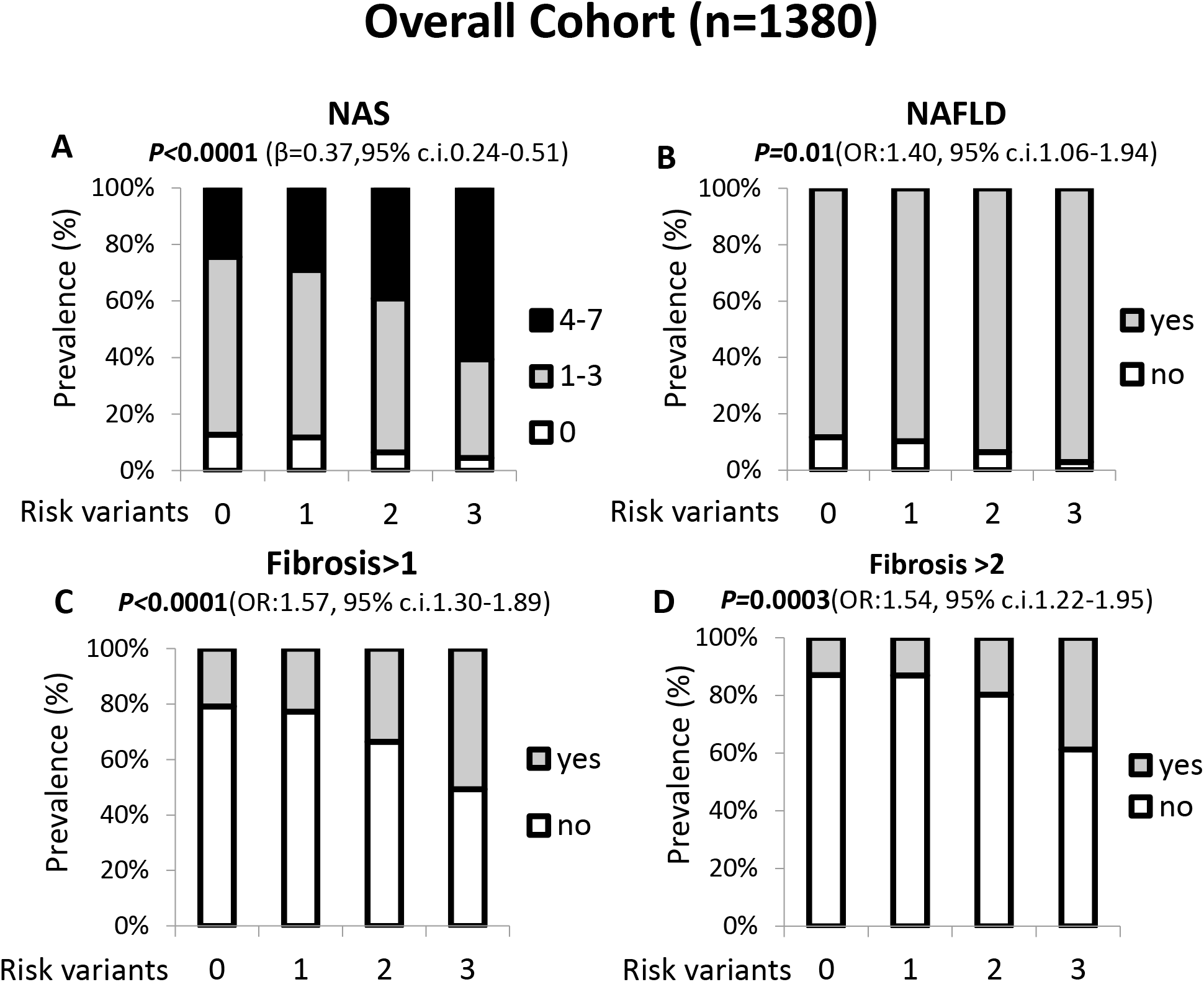

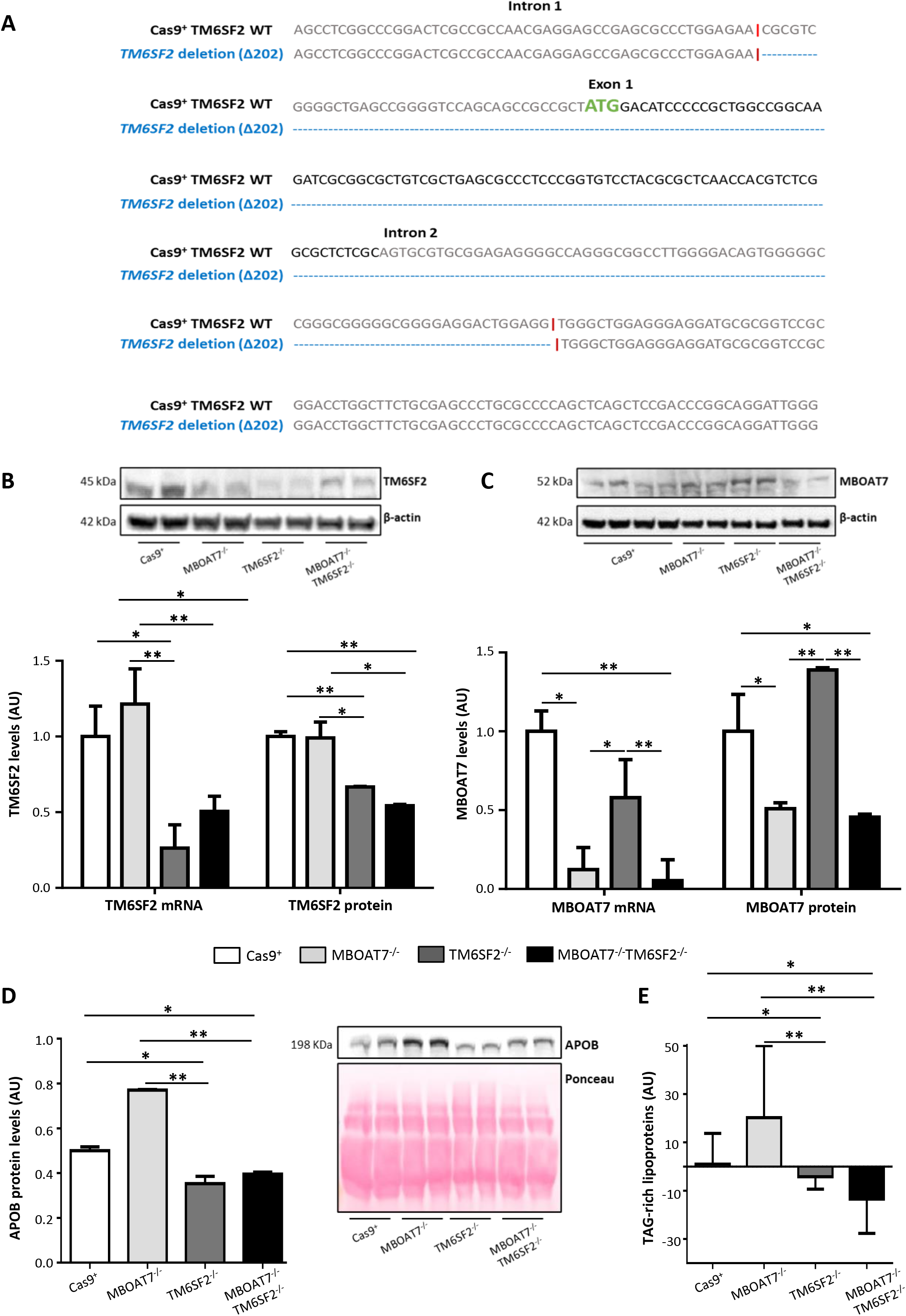

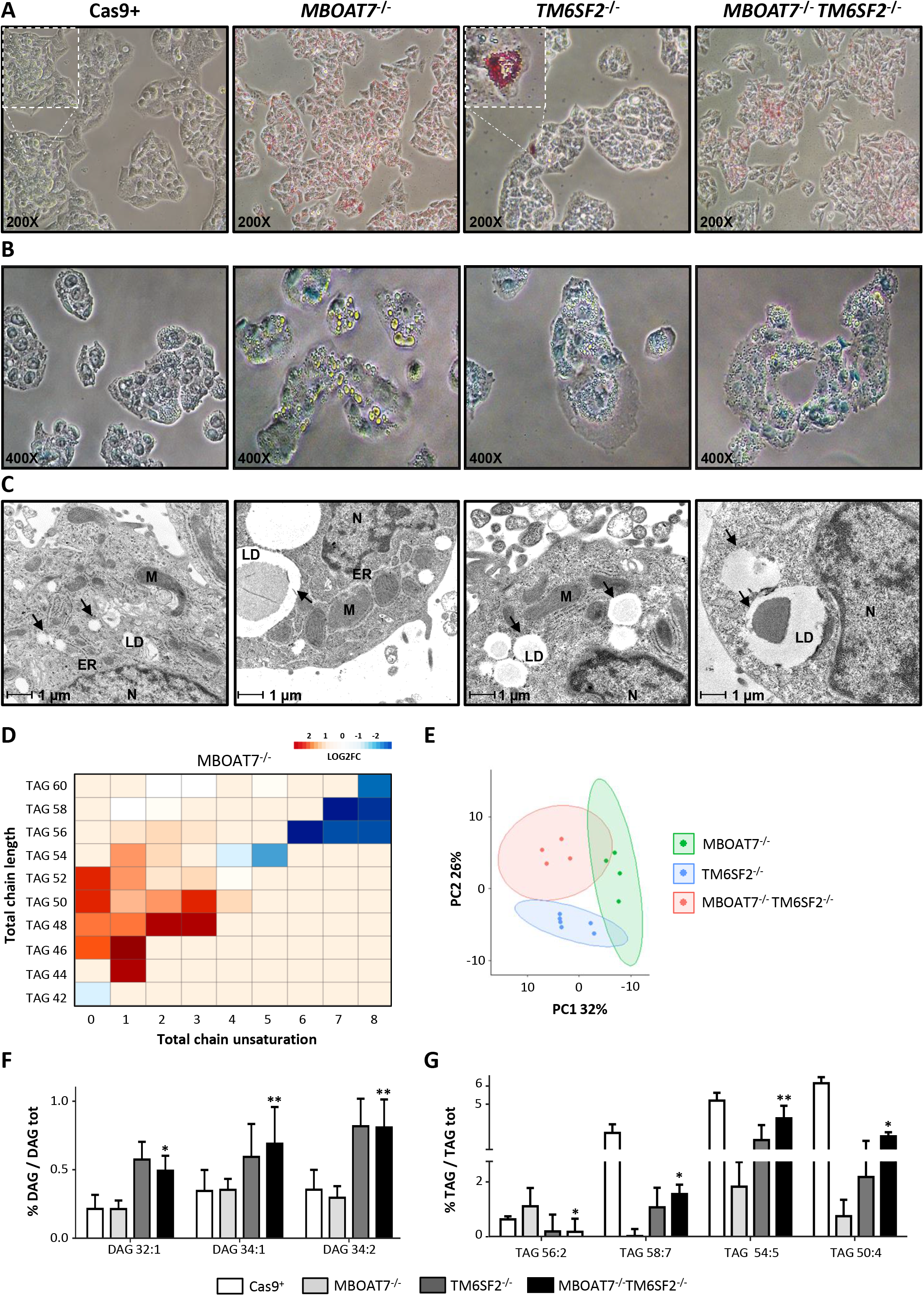

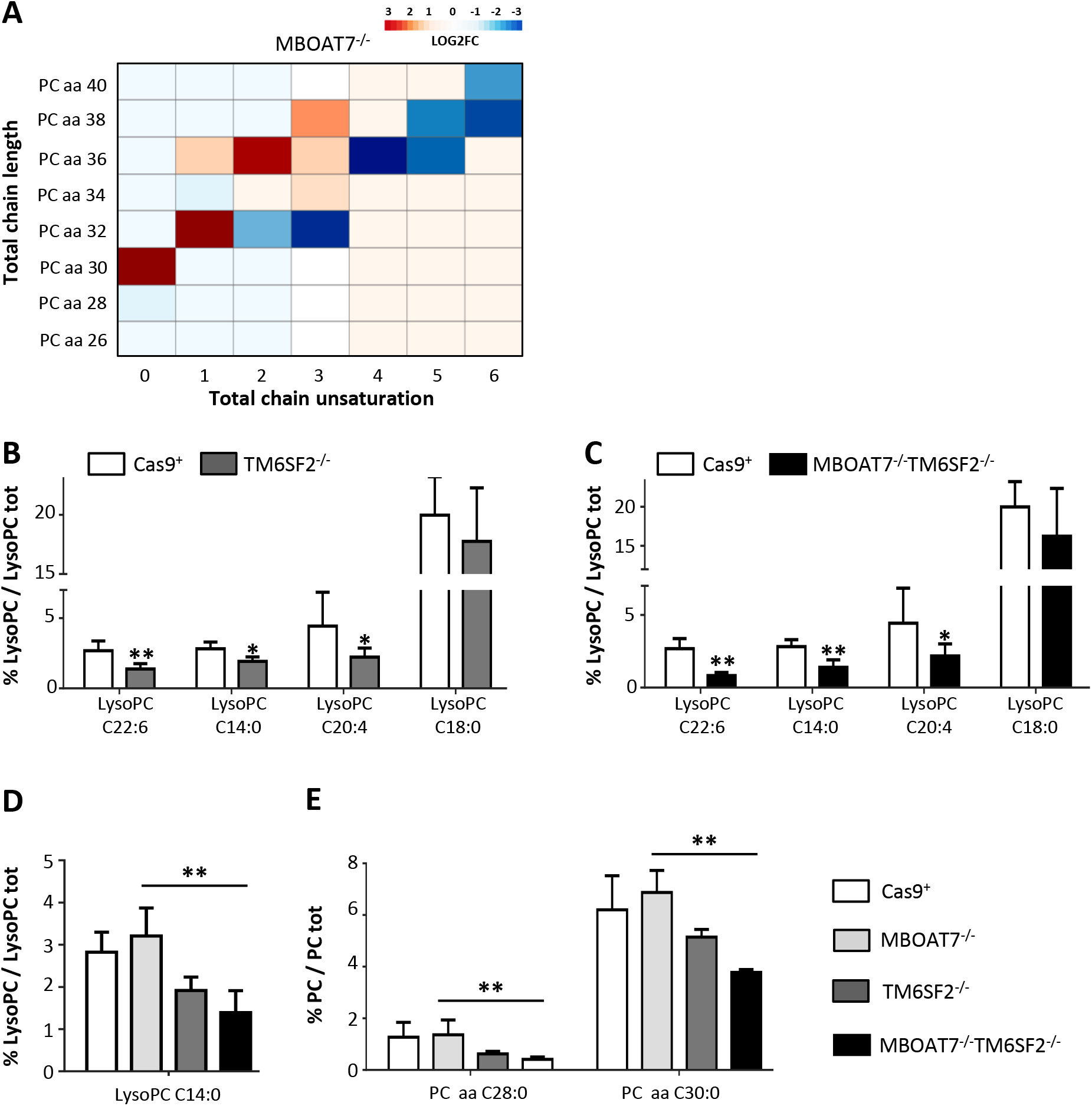

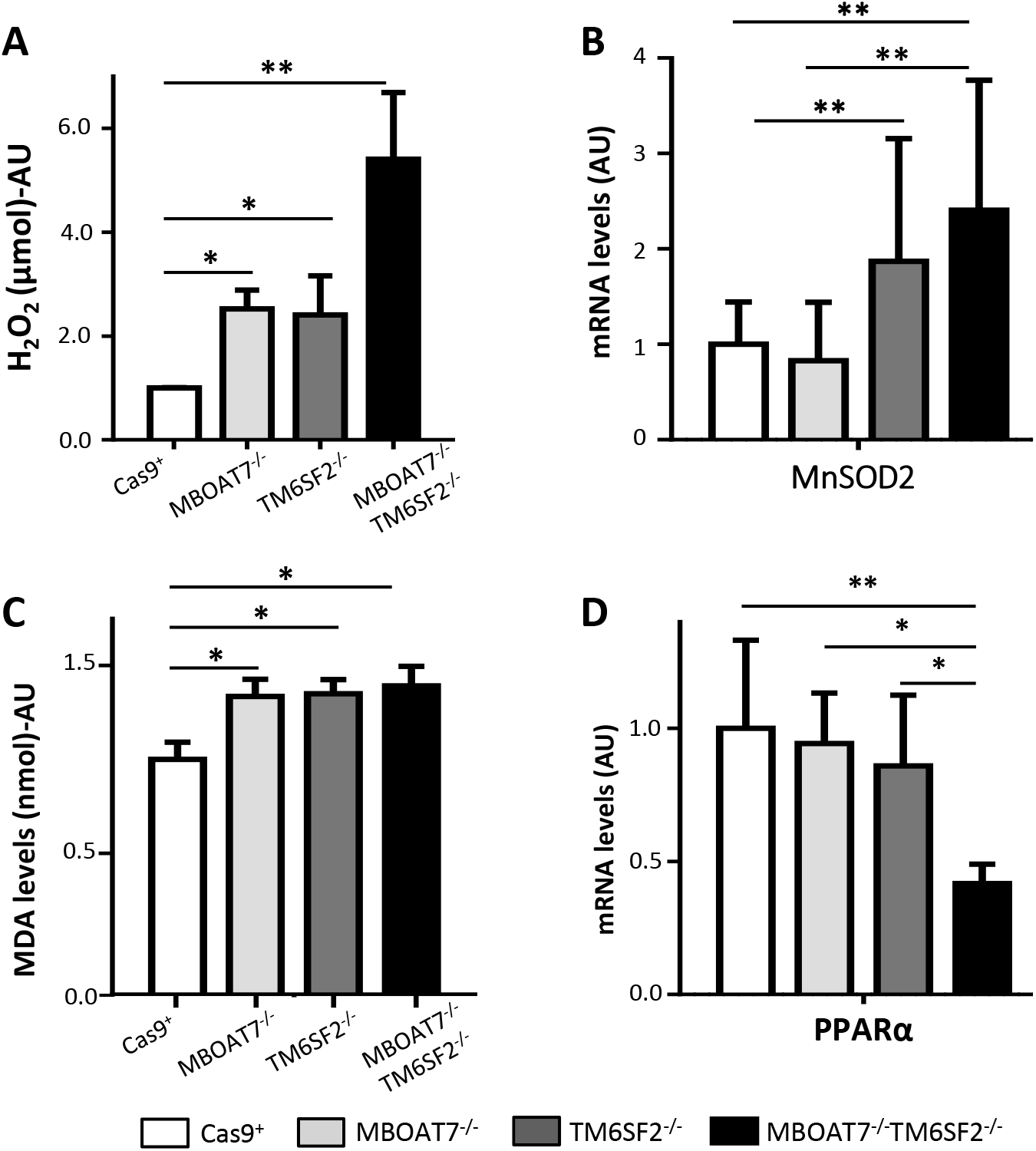

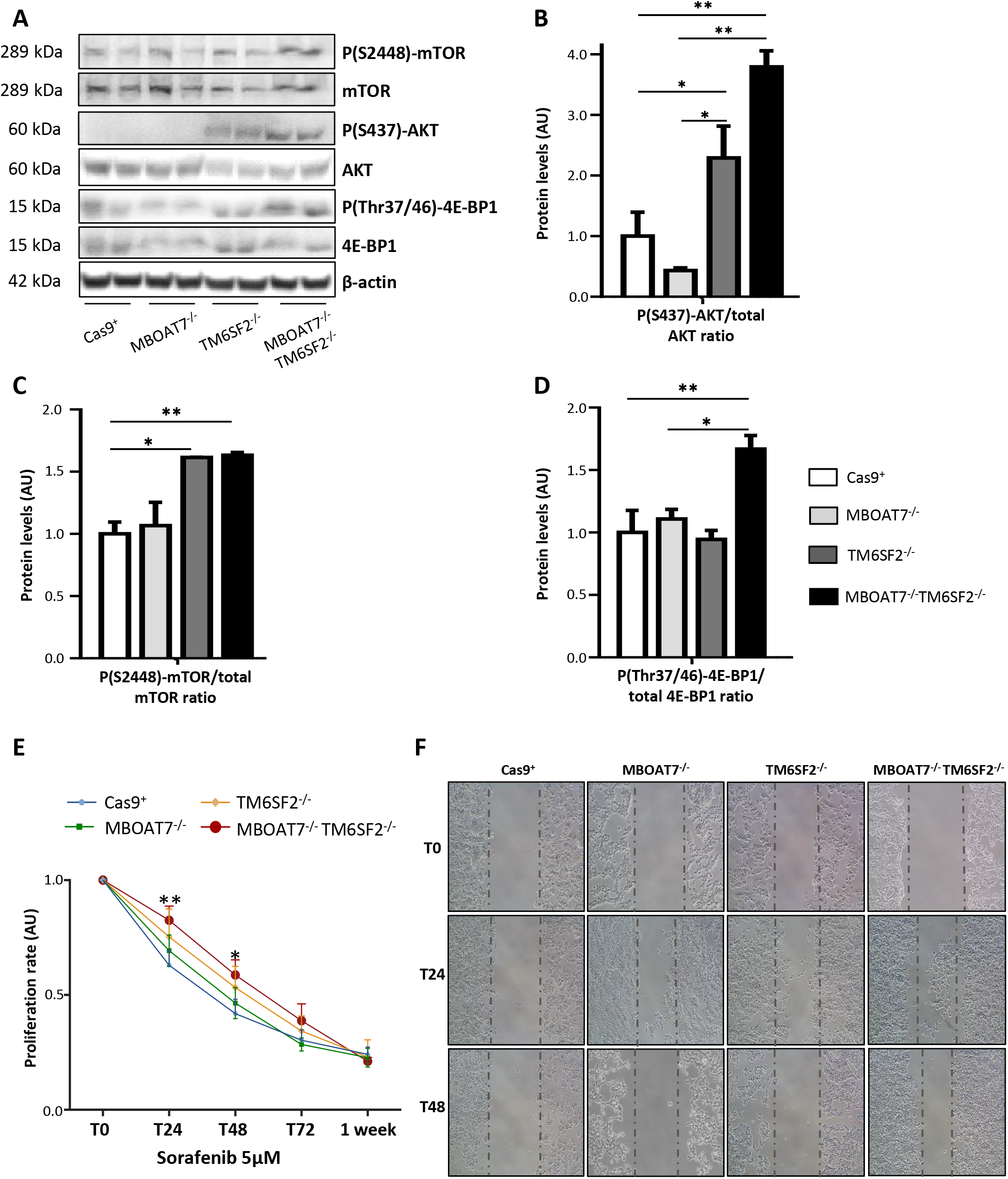

## Notes

**Fundings**: The study was supported by Ricerca Corrente Fondazione IRCCS Cà Granda and Ricerca Finalizzata Ministero della Salute RF-2013-02358319.

### Competing Interest Statement

The authors have declared no competing interest.

