## Supplementary material for "TM6SF2/PNPLA3/MBOAT7 loss-of-function genetic variants impact on NAFLD development and progression both in patients and in *in vitro* models": Suppl materials and Methods

Received: date; Accepted: date; Published: date

### Supplemental materials and methods

##### *Chemicals*

Sorafenib was acquired from St. Cruz Biotechnologies (Santa Cruz, United States). Doxycycline, anti-MBOAT7 and bovine serum albumin (BSA) were purchased from Sigma-Aldrich (St Louis, MO). Anti-P-Ser473 Akt, Akt, P(Thr37/46)-4E-BP1, 4E-BP1, P(Ser2448)-mTOR and mTOR antibodies were acquired from Cell Signaling Technologies (Boston, United States). Anti β-Actin was purchased from Abcam (Cambridge, UK). Anti-Cas9 and anti-PCG1α were obtained from Novus Biologicals (Littleton, U.S.A.). TM6SF2 and APOB-100 antibodies were bought from Life Technologies-ThermoFisher Scientific (Waltham, United States). Dulbecco's modified Eagle's medium (DMEM), fetal bovine serum (FBS), phosphate-buffered saline (PBS), L-Glutamine, penicillin/streptomycin, Trypsin/EDTA, Hank's balanced salt solution (HBSS), fast SYBR green master mix, lipofectamine 3000 transfection reagent, blasticidin, T4 DNA ligase, Fastdigest Eco31I, Ampicillin and DH5α competent cells were obtained Life Technologies-ThermoFisher Scientific (Waltham, United States). PCR Master Mix 2X was bought from BiotechRabbit, (Hennigsdorf, Germany). Clarity western ECL substrate was obtained from Bio-Rad Laboratories (Hercules, United States). CellTiter 96® $AQ_{ueous}$ One Solution Cell Proliferation Assay (MTS) was purchased from Promega Corporation (Fitchburg, U.S.A). The pGL3-U6-sgRNA-PGK-puromycin was a gift from Xingxu Huang (Addgene plasmid #51133; http://n2t.net/addgene:51133; RRID:Addgene_51133; ired from Addgene)[1]. Puromycin and Edit-R Inducible Lentiviral hEF1α-Blast-Cas9 Nuclease Particles were obtained from GE Healthcare Dharmacon Inc[2]. Triacylglycerols (TAGs) Quantification Kit was purchased from BioVision (Milpitas, United States). Cholesterol Colorimetric Assay Kit - HDL and LDL/VLDL, DCF ROS/RNS Colorimetric Assay Kit, Hydrogen Peroxide Assay Kit, Lipid Peroxidation (MDA) Assay Kit (colorimetric/fluorimetric), DNA Damage Assay Kit (A P sites, Colorimetric) and MitoBiogenesis™ In-Cell ELISA Kit were bought from Abcam (Cambridge, UK). VectaMount AQ Mounting Medium was obtained from Maravai Life Sciences Inc. (United Kingdom).

##### *Overall cohort*

The Overall cohort consists of 1380 patients with NAFLD and it has been subdivided in the Hepatology Service cohort (n=1259) and the NAFLD-HCC cohort (n=121).

The Hepatology Service cohort included 1259 unrelated patients of European descent who were consecutively enrolled at the Metabolic Liver Diseases outpatient service (Liver Clinic; n=692) and bariatric surgery center (Bariatric Surgery; n=567) at Fondazione IRCCS Cà Granda, Ospedale Maggiore Policlinico Milano, Milan, Italy. Inclusion criteria were availability of a liver biopsy for suspected NASH or severe obesity, DNA samples, and clinical data. Individuals with excessive alcohol intake (men, >30 g/day; women, >20 g/day), viral and autoimmune hepatitis, or other causes of liver disease were excluded. The study conformed to the Declaration of Helsinki and was approved by the Institutional Review Board of the Fondazione Ca' Granda IRCCS of Milan and relevant Institutions. All participants gave written informed consent.

The NAFLD-Hepatocellular carcinoma (NAFLD-HCC) patients of Italian descent (n=121) were enrolled between January 2008 and January 2015 at the Milan, Udine, Rome hospitals. Diagnosis of HCC was based on the European Association for the Study of the Liver–European Organization for Research and Treatment of Cancer Clinical Practice Guidelines [3]. In the absence of liver biopsy, diagnosis of NAFLD-HCC required detection of ultrasonographic steatosis plus at least one criterion of the metabolic syndrome.

***Histological evaluation***

Steatosis was graded into the following four categories based on the percentage of affected hepatocytes: 0, 0%-4%; 1, 5%-32%; 2, 33%-65%; and 3, 66%-100%. Disease activity was assessed according to the NAFLD activity score, with systematic evaluation of hepatocellular ballooning and necroinflammation; fibrosis was also staged according to the recommendations of the NAFLD Clinical Research Network [4] The scoring of liver biopsies was performed by independent pathologists unaware of patient status and genotype [5, 6]. NASH was diagnosed in the presence of steatosis, lobular necroinflammation, and hepatocellular ballooning.

***Genotyping***

The Overall cohort was genotyped for the rs738409 C>G (PNPLA3 I148M), rs58542926 C>T (TM6SF2 E167K) and rs641738 C>T MBOAT7 risk variants as previously described [10,12]. Genotyping was performed in duplicate using TaqMan 5'- nuclease assays (QuantStudio 3, Thermo Fisher, Waltham, MA). Results of rs738409, rs58542926 and the rs641738 genetic frequencies were compared to those obtained in not-Finnish European healthy individuals included in the 1000 Genome project [7].

***Gene silencing***

*TM6SF2* silencing was induced by exploiting CRISPR/Cas9 genome editing following non-homologous end joining (NHEJ) in both HepG2 hepatoma cells (ATCC-HB-8065, which are homozygous for the I148M PNPLA3 variant, and in HepG2 knockout for the *MBOAT7* gene ($MBOAT7^{-/-}$)[8] already available in our lab, allowing us to obtain a double stable model to study NAFLD. The reduced activity and expression of both MBOAT7 and TM6SF2 lead to an additive risk to develop NAFLD when combined with the I148M PNPLA3 variant [9-13], making the HepG2 cells an ideal model to study *in vitro* the impact MBOAT7 and TM6SF2 loss-of functions coupled to the influence of I148M PNPLA3 on lipid metabolism, hepatocellular damage and carcinogenesis.

HepG2 cells were transfected with a lentiviral vector (Edit-R Inducible Lentiviral hEF1α-Blast-Cas9 Nuclease Particles, GE Healthcare Dharmacon Inc) containing Cas9 gene under the control of doxycycline inducible promoter and blasticidin resistance as selection marker. Concurrently, two single strand DNAs were designed using free online CRISPR Design Tool http://crispr.mit.edu/ (forward: 5'-AAACACGCGCTCAACCACGTCTCG-3'; reverse: 5'- CCGGCGAGACGTGGTTGAGCGCGT) in order to construct a small guide-RNA (sgRNA). Double-stranded oligos were subcloned using T4 DNA ligase (ThermoFisher) into the pGL3-U6-sgRNA-PGK-puromycin plasmid (Addgene plasmid #51133; http://n2t.net/addgene:51133; RRID:Addgene_51133; ired from Addgene)[1] and digested with Fastdigest Eco31I (ThermoFisher). The identity of the small guide-RNA constructs was verified by Sanger sequencing (**Table S4**). The sgRNA was designed to induce TM6SF2 cut by Cas9 transcription star site (ATG) in the first codifying exon of *TM6SF2* gene (isoform 1, transcript variant 1, NM_001001524.3) to induce TM6SF2 knockout altering its mRNA transcription. After doxycycline-induced Cas9 expression, HepG2/Cas9 positive clones ($Cas9^+$) were transfected (Lipofectamine 3000 transfection reagent, Thermofisher Scientific) with pGL3-U6-sgRNA-PGK-puromycin plasmid containing the sgRNA under the control of cytomegalovirus promoter (U6), which was previously subcloned in Dh5α competent cells. Positive clones carrying the *TM6SF2* mutations were selected exploiting puromycin resistance gene as selection marker (GE healthcare Dharmacon Inc). After single-cell clonal expansion and single clone-derived colonies formation generated by the limiting dilution, efficiency of gene editing was tested by Sanger sequencing (about 100 clones *per* condition have been screened). Potential off-target modifications were checked in the genome region of interest. In both $CAS9^+$ and $MBOAT7^{-/-}$ cells, we identified the same *TM6SF2* deletion of 202 nucleotides (Δ202) in homozygosis, referred to as $TM6SF2^{-/-}$ and $MBOAT7^{-/-}$ $TM6SF2^{-/-}$, respectively. $Cas9^+$ cells wild type (Wt) in the *TM6SF2* gene were used as control group.

***Transmission Electron Microscopy (TEM)***

$Cas9^+$, $MBOAT7^{-/-}$, $TM6SF2^{-/-}$ and $MBOAT7^{-/-}TM6SF2^{-/}$ cells were cultured as monolayer (70-80% confluent) and trypsinized to obtain cell suspension. Cells were fixed with aldehyde mixture (4% Paraformaldehyde + Glutaraldehyde 2.5% in cacodylate buffer: pH 7.4) at 4°C overnight. After primary fixation, cells were washed repeatedly with cacodylate buffer and were post-fixed in 1% osmium tetroxide ($OsO_4$) for 2 hours in dark. Next, samples were left in 1.5% potassium ferrocyanide dissolved in 0.1 M cacodylate for 1 hour on ice. Each sample was stained with 0.5% 5-uranyl acetate in water overnight. Finally, samples were dehydrated with rising ethanol series, embedded in an Epon resin, and polymerized in an oven at 60°C for 48 hours. Ultrathin (70-90 nm) sections were collected on nickel grids and observed with a ZEISS Leo 912 AB Omega TEM [14].

***In vitro treatments***

When specified, $Cas9^+$, $MBOAT7^{-/-}$, $TM6SF2^{-/-}$ and $MBOAT7^{-/-}$ $TM6SF2^{-/-}$ were treated with either Sorafenib (Santa Cruz, United States) at final concentration of 5 µM or dimethyl-sulfoxide (DMSO) vehicle for 24-48-72 hours and 1 week. Treatments were freshly prepared and administered daily.

***DNA extraction and Sanger sequencing***

DNA was extracted using phenol/chloroform method. Next, DNA concentration and quality were assessed by Nanodrop 1000 microvolume 42 spectrophotometer (ThermoFisher Scientific, U.S.A.). 1 µL of DNA was used to verify single guide RNA (sgRNA) insertion in the pGL3-U6-sgRNA-PGK-puromycin plasmid [1] and to detect *TM6SF2* mutations in the exon I of *TM6SF2* sequence. PCR products were run on agarose gel 3% electrophoresis, purified with a vacuum pump, and used to perform the Sanger sequencing protocol (Big Dye mix, Applied Biosystem). Primers used for amplification and sequencing are listed in **Table S5**.

***Gene expression analysis***

RNA was extracted from cell cultures using Trizol reagent (Life Technologies-ThermoFisher Scientific, Carlsbad, U.S.A). 1µg of total RNA was retro-transcribed with VILO random hexamers synthesis system (Life Technologies-ThermoFisher Scientific, Carlsbad, U.S.A). Quantitative real time PCR (qRT-PCR) was performed by an ABI 7500 fast thermocycler (Life Technologies), using the TaqMan Universal PCR Master Mix (Life Technologies, Carlsbad, CA) and TaqMan probes for human MBOAT7, TM6SF2, glucokinase (GCK), phosphofructokinase (PFK), glyceraldehyde-3-phosphate (G3P) and GAPDH. The SYBR Green chemistry (Fast SYBR Green Master Mix; Life Technologies) was used for all other genes. All reactions were delivered in triplicate. Data were normalized to the GAPDH or β-actin gene expression and results were expressed as arbitrary units (AU) or fold increase as indicated in bar graphs. Primers are listed in **Table S5-S6.**

***Western Blot Analysis***

Total protein lysates were extracted from 2 cell cultures, using RIPA buffer containing 1 mmol/L Na-orthovanadate, 200 mmol/L phenylmethyl sulfonyl fluoride and 0.02 μg/μL aprotinin. Samples were pooled prior electrophoretic separation and all reactions were performed in duplicate. Then equal amounts of proteins (50 μg) were separated by SDS-PAGE, transferred electrophoretically to nitrocellulose membrane (BioRad, Hercules, CA) and incubated with specific antibodies overnight. At least, three independent lots of freshly extracted proteins were used for experiments. Antibodies and concentration used are listed in **Table S7**.

***Oil Red O (ORO) staining***

Cells were plated on 6-wells plate ($5x10^5$ cells/well) in duplicate and left overnight in DMEM medium containing 10% FBS, 1% L-glutamine and 1% Penicillin/Streptomycin. After 24 hours, grow media was removed and cells were kept 24h in quiescent medium, containing 0.5% BSA, 1% L-glutamine and 1% Penicillin/Streptomycin. The day after, we performed Oil Red O (ORO) staining, which is a soluble red powder with high affinity for neutral TAGs and lipids stored in the lipid droplets (LDs). Quiescent medium was removed, and the 6-well plates were gently rinsed with 2 mL of sterile PBS 1X. Next, cells were fixed with 4% formalin for 1h at room temperature. After fixation, each sample was washed with sterile water and 60% isopropanol was added for 5 minutes. Concurrently, we prepared ORO working solution by mixing 3 parts of ORO stock solution (300 mg of Red Oil powder in 100 mL di isopropanol 100%) and 2 parts of sterile water, following filtration. 1 mL/well of ORO working solution was added to each sample and left for 40 minutes. Finally, plates were rinsed with tap water, paying attention not to disrupt the monolayer. Lipid droplets content will be visualized in pink-red color. ORO positive area will be quantified by ImageJ software in 10 random micrographs (magnification 200x) by calculating the ORO positive area as percentage of pixels above the threshold value with respect to the total pixels per area.

***Immunocytochemistry (ICC)***

$1x10^6$ cells were seeded on coverslip lodged in a 6-well plate in duplicate and kept overnight in DMEM medium containing 10% FBS, 1% L-glutamine and 1% Penicillin/Streptomycin. Next, hepatocytes were fixed in 4% formalin for 1h and permeabilized in 0.3% Triton-X 100. Cells were incubated in 5% BSA for 30 minutes and with anti-PGC1α primary antibody overnight at 4°C. Then, each sample was incubated with anti-rabbit HRP-conjugated antibody and 3,3′-Diaminobenzidine (DAB) was provided as chromogen. Nucleus were counterstained with hematoxylin. Finally, samples were mounted with a drop aqueous VectaMount AQ Mounting Medium (Maravai Life Sciences Inc. United Kingdom).

***Evaluation of MT-COX1 expression***

Mitobiogenesis In-Cell ELISA Kit Abcam (Cambridge, UK) exploits a quantitative immunocytochemistry to measure mitochondrially-encoded COX-I (MT-COX1) and nuclear -encoded SDHA protein levels in cultured cells and it was performed following the manufacturer' instructions[15]. Briefly, cells are seeded in a 96-well plate ($3x10^5$/well) in triplicate, fixed with 4% paraformaldehyde and permeabilized through 1X Triton-X 100. Targets of interest are detected with highly specific, well-characterized cocktail of monoclonal antibodies, which were incubated overnight at 4°C. Then, AP-labelled and HRP-labeled secondary antibodies are used to generate a colorimetric reaction that could be measured at 405 and 600 nm, respectively.

***Lipidomic analysis***

For lipidomic analysis, lipid classes were separated with ultra-high pressure liquid chromatography (UHPLC) equipped with ZORBAX Eclipse Plus C18 2.1x100mm 1.8 µm columns (Agilent, Santa Clara CA), and lipids concentrations were measured by MS-QTOF (Agilent UHPLC 1290 Infinity-6540 QTOF, Santa Clara, CA) in both positive (for ceramides (Cer), sphingomyelins (SMs), phosphatidylethanolamines (PEs), Lyso-PEs, phosphatidylcholines (PCs), Lyso-PCs, diacylglycerols (DAGs), triacylglycerols (TAG)) and negative electrospray ionization for phosphatidylinositols (PIs). Mobile phase A consists in water with 0.1% formic acid in positive mode and 5 mM Ammonium Acetate in negative mode; mobile phase B consists in acetonitrile: isopropanol (1:1) with 0.1% formic acid in positive and 5 mM Ammonium Acetate in negative mode. Analysis of the lipid profiles was performed by Mass Hunter Profinder software coupled with PCDL Manager (Agilent, Santa Clara, CA) that allows to extract, identify and quantify selected lipid compounds on the bases of the accurate mass m/z, retention time and ion abundance. The software calculates the area under the curve of identified ions. For the lipid species, ions are extracted as adducts [M + H] + and [M + Na] + when samples were run in positive mode and as [M-H]- and [M + HAc-H]- when samples were run in negative mode (HAc = CH3COO-). All sets of samples were run together in order to avoid batch differences. Standard Reference Materia SRM 1950 (National Institute of Science and Technology NIST, US) was run as quality control during each run (positive and negative).

Lipids measured in positive mode concentrations were quantified using internal standards, i.e. SM (d18:1/17:0), Ceramide (d18:1/17:0), PE (17:0/17:0), PC (17:0/17:0), Lyso-PC 17:0, DG (17:0/17:0) and TAG (15:/15:0/15:0 (Avanti Polar Lipids, Alabaster, AL and Larodan, Solna, SE). PIs and Lyso-PIs were quantified using as internal standards either Phosphatidylglycerol (PG)34:0 or ceramide (d18:1/17:0), which are visible in negative mode and have a retention time close to PI and Lyso-PI. Both internal standards gave similar concentration values for PIs and Lyso-PIs.

Qualitative analyses were shown in figures. Data were analyzed as log2 fold change (log2FC) ratio considering chain length and number of double bonds (*i.e.* degree of unsaturation). Principal Component Analysis (**Figure S3E**) was performed on the log2 fold change ratios between $MBOAT7^{-/-}$, $TM6SF2^{-/-}$, $MBOAT7^{-/-}TM6SF2^{-/-}$ and Cas9 + lipidomic data, using R package FactoMineR[16] (1). Since PCA is an unsupervised method based on a linear combination of features, it allows to evaluate the different changes induced by the mutations on the whole lipidomic profile, even with a small number of samples.

***Evaluation of reactive oxygen species (ROS) and lipid peroxidation***

Cas9$^{+}$, MBOAT7$^{-/-}$, TM6SF2$^{-/-}$ and MBOAT7$^{-/-}$TM6SF2$^{-/}$ cells were cultured as monolayer. At 70-80% of confluence, they were stained with MitoSOX Red (1.25 µM; Invitrogen) for 30 minutes at 37°C, and subsequently harvested for FACS analysis (excitation 488 nm, emission 690/50 nm; Acea Bioscience, USA). Increased red fluorescence was correlated with ROS formation. Data were collected from at least 20000 cells and six replicates per condition. Lipid peroxidation was analyzed by staining cells with BODIPY 581/591 C11 (2µM; Thermo Fisher Scientific, Darmstadt, Germany) for 1 hour at 37°C. Cells were then harvested for FACS analysis (excitation 488nm, emission 525/30nm and 585/50nm) and the shift from red to green fluorescence was used to evaluate lipid peroxidation. Data were collected from at least 20,000 cells and six replicates per condition.

***Evaluation of Complex I enzymatic activity***

$10^6$ cells were resuspended in proper buffer (pH 7.2). Protein extraction was performed by sonicating cell pellets at 50 W (10 s) for 3 times. Lysates were centrifuged at 750g for 10 min and supernatant was recovered. Lowry method was used for protein quantification. A Lambda 2 Parkin Elmer spectrophotometer was used to assess enzymatic activities. Analyses were performed at specific wavelengths for each enzymatic activity after preparing proper solutions as previously described[17, 18]. NADH dehydrogenase (340 nm): H2O (455µl), K-phosphate pH 7.5 0.1 M (340µl), K3[Fe(CN)6] 17 mM (100µl), NADH2 mM (100µl), homogenate (5µl). NADH ubiquinone 1 reductase (340 nm): H2O (610µl), K-phosphate pH 7.5 0.1 M (200µl), albumin1% (100µl), NADH 2 mM (70µl), homogenate (10µl), sodium azide100 mM (10µl), CoQ1 6 mM (5µl); activity measured after rotenone administration (1 mM, 5µl) was subtracted. Citrate synthase (412 nm): H2O (800µl), DTNB 1 mM (100µl), oxaloacetic acid 10 mM (50µl), Acetyl-CoA 10 mM (30µl), homogenate (20µl). Experiments were performed at 30 °C. Analyses were performed through a Parkin Elmer software. Measurements were normalized over the activity level of citrate synthase, a stable matrix mitochondrial enzyme; this latter step was per-formed in order to normalize respiratory chain activity over mitochondrial mass.2.6.

***Measurement of ATP rate***

The ATP (adenosine triphosphate) production in live cells was measured by Agilent Seahorse XF Real-Time ATP Rate Assay (Agilent Technologies, USA), which quantifies the rate of ATP from glycolysis and mitochondria in real time. The assay employs a sequential injection of specific mitochondrial activators and inhibitors and reports multiple parameters, including glycolytic ATP production rate, mitochondrial ATP production rate, total ATP production rate and XF ATP rate index. Briefly, cells were plated in Seahorse XF24-well microplate (40,000 cells per well, Agilent) and the day of the assay the cell culture growth medium was replaced by the assay medium, supplemented of glucose, pyruvate and glutamine, and cells incubated at 37°C for 60 minutes. Oxygen Consumption Rate (OCR) and Extracellular Acidification Rate (ECAR) were measured using XFe24 Analyzer, with three baseline measurements recorded before and after adding Oligomycin (ATP synthase inhibitor; 1.5 µM) and a mix of Rotenone and Antimycin A (complex I and III inhibitors, respectively; 0.5 µM).

***Cell proliferation assay***

$Cas9^{+}$, $MBOAT7^{-/-}$, $TM6SF2^{-/-}$ and $MBOAT7^{-/-}TM6SF2^{-/-}$ ($2.5*10^{2}$ cells/well) were seeded in in 96-well plate in quadruplicate and incubated under normal culture conditions overnight. Cell proliferation was measured at both baseline and upon Sorafenib exposure using the CellTiter96-Aqueous One Solution Cell Proliferation Assay (MTS) kit (Promega Corporation, Fitchburg, USA) according to the manufacturer's instructions. A dose-response curve was performed by testing 1-2.5-5-10-50 µM of Sorafenib treatments to determine the minimum concentration required to evaluate cell viability. Administration of Sorafenib at very low concentrations (1 and 2.5 µM) resulted inefficacy to modulate cell survival, whereas the higher ones (10 and 50 µM) showed high mortality rate. Therefore, after a day, cells were treated with either Sorafenib at the final concentration of 5µM or vehicle (DMSO), which is further consistent with the current literature[19, 20].

Freshly growth media with or without Sorafenib was provided for 24-48-72 h and 1 week. MTS reagent (20 µL/well) was added to the cells followed by incubation for 4 h in 5% $CO_2$ humidified incubator at 37 °C, and the absorbance was measured at 490 nm, at 0, 24, 48-74 h and after 1 week. At least three independent experiments were carried out.

***Wound healing assay***

$Cas9^{+}$, $MBOAT7^{-/-}$, $TM6SF2^{-/-}$ and $MBOAT7^{-/-}TM6SF2^{-/}$ models were plated on 6-well plate ($8x10^{5}$/well) and incubated with DMEM medium containing 10% FBS, 1% L-glutamine and 1% Penicillin/Streptomycin. A fine scratch was introduced using a sterile pipette tip in a monolayer of cells at ~90% confluency. Cells were then treated with sorafenib at the final concentration of 5 µM or vehicle (DMSO) for 24 and 48 h. The wounds were photographed (100X objective) at 24 h and 48 h. Each experiment was performed in triplicate.

### Supplemental results

***CRISPR/Cas9-mediated gene editing in hepatocytes to model NAFLD***

To explore whether *TM6SF2* and *MBOAT7* loss-of functions in the context of I148M PNPLA3 genetic background may exert an additive effect in hepatocytes, in terms of fat accumulation, lipid metabolism, hepatocellular stress and carcinogenesis, we exploited CRISPR/Cas9 technology to induce genetic deficiency of the *MBOAT7* (MBOAT7$^{-/-}$)[8], *TM6SF2* (TM6SF2$^{-/-}$) or both (MBOAT7$^{-/-}$ TM6SF2$^{-/-}$) in HepG2 cells. Sanger sequencing confirmed *TM6SF2* silencing due to a deletion of 202 nucleotides (Δ202) cutting the ATG site in both TM6SF2$^{-/-}$ and MBOAT7$^{-/-}$ TM6SF2$^{-/-}$ clones compared to the Wt reference sequence (**Figure S2A**). As expected, TM6SF2 mRNA and protein levels were reduced in TM6SF2$^{-/-}$ and MBOAT7$^{-/-}$ TM6SF2$^{-/-}$ cells compared to Cas9$^{+}$ and MBOAT7$^{-/-}$ ones ($p<0.0001$ at ANOVA; adjusted $p<0.05$ *vs* Cas9$^{+}$ and MBOAT7$^{-/-}$, **Figure S2B**). Likewise, the MBOAT7 expression was lower only in MBOAT7$^{-/-}$ and MBOAT7$^{-/-}$ TM6SF2$^{-/-}$ cells ($p=0.0002$ at ANOVA; adjusted $p<0.05$ *vs* Cas9$^{+}$ and TM6SF2$^{-/-}$, **Figure S2C**), thus recapitulating the human condition of genetic NAFLD in which the rs641738 and the E167K variants cause a reduction of MBOAT7 and TM6SF2 levels, respectively.

To assessed whether *TM6SF2* deletion impacted on its functional role, we evaluated both ApoB protein levels and TAG-rich lipoproteins' export in cell supernatants. MBOAT7$^{-/-}$ cells highly enhanced the ApoB and TAG-rich lipoproteins secretion, probably as a compensatory mechanism to remove intracellular lipids (**Figure S2D-E**). Conversely, either TM6SF2$^{-/-}$ and MBOAT7$^{-/-}$ TM6SF2$^{-/-}$ reduced ApoB levels (adjusted $p<0.05$ *vs* Cas9$^{+}$ and $p<0.01$ *vs* MBOAT7$^{-/-}$, **Figure S2D**) and completely abrogated TAG-rich lipoproteins' release compared to both Cas9$^{+}$ and MBOAT7$^{-/-}$ cells ($p=0.0005$ at ANOVA, adjusted $p<0.05$ *vs* control and $p<0.01$ *vs* MBOAT7$^{-/-}$, respectively, **Figure S2E**) thereby supporting that lipoproteins synthesis and export was affected by *TM6SF2* silencing.

***Synergic contribution of the MBOAT7 and TM6SF2 silencing to Sorafenib response***

We investigated whether the co-presence of Cas9-induced mutations in HepG2 cells may also affect the pharmacological response to Sorafenib, a multikinase inhibitor approved for the treatment of advanced HCC. Therefore, we exposed Cas9$^{+}$, MBOAT7$^{-/-}$, TM6SF2$^{-/-}$ and MBOAT7$^{-/-}$ TM6SF2$^{-/-}$ models either to Sorafenib (5µM) or vehicle for 24-48-72 hours and 1 week. While just after 24 hours the proliferation rate of Cas9$^{+}$ was remarkably reduced at MTS assay, MBOAT7$^{-/-}$ or TM6SF2$^{-/-}$ cells delayed cell death in response to Sorafenib (adjusted $p<0.05$ *vs* Cas9$^{+}$, **Figure S6E**) and survival was even higher in clones bearing both mutations (adjusted $p<0.01$ *vs* Cas9$^{+}$, **Figure S6E**). In addition, the MBOAT7$^{-/-}$ TM6SF2$^{-/-}$ cells showed a significant resistance to Sorafenib cytotoxicity until 48 hours (adjusted $p<0.05$ *vs* Cas9$^{+}$, **Figure S6E**). Similarly, after 24 hours of Sorafenib (5µM) exposure, the MBOAT7$^{-/-}$, TM6SF2$^{-/-}$ and MBOAT7$^{-/-}$ TM6SF2$^{-/-}$ cells were able to migrate from one side to the other side of the wound healing in order to repair the scratch area and the major effect was observed in the MBOAT7$^{-/-}$ TM6SF2$^{-/-}$ model (**Figure S6F**).

#### Tables

**Table S1.** Demographic, anthropometric, and clinical features of the Overall Cohort (n=1380), including the Hepatology Service Cohort (n=1259) and the NAFLD-HCC (n=121), stratified for enrollment criteria.

| | **Overall cohort (n=1380)** | **Hepatology Service Cohort (n=1259)** | **NAFLD-HCC (n=121)** | ***P*-value*** |
|---|---|---|---|---|
| Sex, M | 739 (53.5) | 650 (51.6) | 89 (73.5) | 0.06 |
| Age, years | 50.3±13.5 | 47.86±12.6 | 67.64±10.0 | **<0.0001** |
| BMI, kg/m2 | 34.3±8.7 | 34.7±8.8 | 28.7±5.14 | **0.01** |
| IFG/T2D, yes | 369 (26.7) | 293 (23.3) | 76 (62.8) | **0.005** |
| HOMA-IR | 5.26±7.96 | 5.01±5.49 | 12.4±31.3 | **<0.0001** |
| Insulin, IU/ml | 20.9±24.0 | 20.3±16.9 | 39.4±91.6 | **0.0003** |
| Total cholesterol, mmol/L | 5.14±1.07 | 5.18±1.04 | 4.28 ±.1.18 | **<0.0001** |
| LDL cholesterol, mmol/L | 3.15±0.97 | 3.19±0.95 | 2.4±0.99 | **<0.0001** |
| HDL cholesterol, mmol/L | 1.29±0.38 | 1.29±0.37 | 1.31±0.51 | 0.44 |
| Triglycerides, mmol/L | 1.62±1.00 | 1.63±0.94 | 1.41±1.81 | **0.03** |
| ALT, IU/l | 4.49 {2.99-4.04} | 3.46{2.99-4.04} | 3.61 {3.29-3.98} | 0.77 |
| AST, IU/l | 3.21 {2.94-3.61} | 3.21{2.89-3.58} | 3.62 {3.22-4.04} | **0.0074** |
| **Number of Risk variants** | | | | ***P*-value**** |
| 0 | 172 (12.46) | 164 (13.02) | 6 (4.9) | **0.009** |
| 1 | 574 (42.03) | 538 (42.73) | 38 (31.4) | **0.01** |
| 2 | 552 (40) | 489 (38.84) | 63 (52) | **0.004** |
| 3 | 82 (5.94) | 68 (5.4) | 14 (11.5) | **0.006** |

Values are reported as mean ±SD, number (%) or median {IQR}, as appropriate. BMI: body mass index; IFG: impaired fasting glucose; T2D: type 2 diabetes. Characteristics of participants were compared across class enrollment criteria using linear regression model (for continuous variables) or logistic regression model (for categorial characteristics).*Models were adjusted for gender, age, BMI, IFG/T2D, class enrollment and number of 3 at-risk variants (I148M PNPLA3, E167K TM6SF2 and the rs641738 C>T *MBOAT7*). **The frequencies for each risk variants subgroup were compared using chi-squared ($\chi^2$) test. p<0.05 was considered statistically significant. 0: indicates the absence of risk variants; 1-2-3 indicates the total number of risk variants carried.

*Hepatology service cohort *vs* NAFLD-HCC cohort

**Table S2.** Demographic, anthropometric, and clinical features of Overall cohort (n=1380) stratified for number of *PNPLA3* I148M, *MBOAT7* rs641738 and *TM6SF2* E167K risk variants

| | **Number of risk variants** | | | | |
|---|---|---|---|---|---|
| | **0 (n=172)** | **1 (n=574)** | **2 (n=552)** | **3 (n=82)** | ***P*-value*** |
| Sex, M | 78 (45.9) | 304 (52.9) | 305 (55.3) | 51 (62.2) | 0.27 |
| Age, years | 49.02±12.17 | 48.69±13.17 | 49.71±13.96 | 52.33±14.70 | 0.80 |
| BMI, kg/m2 | 35.77±8.37 | 34.15±8.68 | 34.48±8.85 | 31.71±8.26 | **0.02** |
| IFG/T2D, yes | 42 (24.41) | 138 (24.04) | 159 (28.8) | 29 (35.36) | 0.10 |
| HOMA-IR | 4.34±3.30 | 4.98±3.82 | 5.81±11.57 | 5.25±3.92 | 0.16 |
| Insulin, IU/ml | 18.7±13.03 | 20.08±13.96 | 22.63±34.01 | 20.50±11.3 | 0.16 |
| Total cholesterol, mmol/L | 5.21±1.09 | 5.24±1.07 | 5.07±1.08 | 4.77±0.80 | **0.006** |
| LDL cholesterol, mmol/L | 3.23±0.94 | 3.20±0.98 | 3.13±0.99 | 2.87±0.70 | 0.06 |
| HDL cholesterol, mmol/L | 1.36±0.37 | 1.33±0.41 | 1.25±0.34 | 1.22±0.35 | **0.001** |
| Triglycerides, mmol/L | 1.51±0.96 | 1.69±1.07 | 1.59±0.95 | 1.54±1.02 | 0.57 |
| ALT, IU/l | 3.33 {2.91-3.97} | 3.40 {2.94-3.97} | 3.58 {3.07-4.13} | 3.67 {3.25-4.14} | **0.003** |
| AST, IU/l | 3.46 {3.17-3.79} | 3.17{2.89-3.55} | 3.29 {2.99-3.66} | 3.46 {3.17-3.79} | **<0.0001** |

Values are reported as mean ± SD, number (%) or median {IQR}, as appropriate. BMI: body mass index; IFG: impaired fasting glucose; T2D: type 2 diabetes. Characteristics of participants were compared across the increasing number of at-risk variants (I148M *PNPLA3*, E167K *TM6SF2* and the rs641738 in *TMC4/MBOAT7* locus) using linear regression model (for continuous variables) or logistic regression model (for categorial characteristics). *Models were adjusted for gender, age, BMI, IFG/T2D and number of 3 at-risk variants. 0: indicates the absence of risk variants; 1-2-3 indicates the total number of risk variants carried.

**Table S3.** Diameter, circumference, and areas of LDs evaluated by TEM analysis and stratified according to the genetic background of the HepG2 cells

| | **Diameter (d)** | **ANOVA** | **Student t-test** | ***P-value**** |
|---|---|---|---|---|
| Cas$^{+}$ | 0.38 {0.33-0.43} | <0.0001 | | |
| MBOAT7$^{-/-}$ | 2.88 {2.59-3.18} | <0.0001 | <0.0001 | **<0.0001** |
| TM6SF2$^{-/-}$ | 1.05 {0.91-1.16} | <0.0001 | <0.0001 | **<0.0001** |
| MBOAT7$^{-/-}$ TM6SF2$^{-/-}$ | 2.42 {2.04-2.72} | <0.0001 | <0.0001 | **<0.0001** |
| | **Circumference (µm)** | **ANOVA** | **Student t-test** | ***P-value**** |
| Cas$^{+}$ | 1.19 {1.05-1.35} | <0.0001 | | |
| MBOAT7$^{-/-}$ | 9.09 {8.13-10.0} | <0.0001 | <0.0001 | **<0.0001** |
| TM6SF2$^{-/-}$ | 3.31 {2.87-3.65} | <0.0001 | <0.0001 | **<0.0001** |
| MBOAT7$^{-/-}$ TM6SF2$^{-/-}$ | 7.60 {6.42-8.55} | <0.0001 | <0.0001 | **<0.0001** |
| | **Area (µm$^{2}$)** | **ANOVA** | **Student t-test** | ***P-value**** |
| Cas$^{+}$ | 0.11 {0.09-0.14} | <0.0001 | | |
| MBOAT7$^{-/-}$ | 6.51 {5.26-7.98} | <0.0001 | <0.0001 | **<0.0001** |
| TM6SF2$^{-/-}$ | 0.87{0.65-1.06} | <0.0001 | 0.003 | **<0.0001** |
| MBOAT7$^{-/-}$ TM6SF2$^{-/-}$ | 4.60 {3.28-5.82} | <0.0001 | <0.0001 | **<0.0001** |

Values are reported as median {IQR}. **P*-values are adjusted at *post hoc* Dunn's multiple comparison test and compared to Cas9$^{+}$ control group.

**Table S4.** ER cisternae width calculated from TEM micrographs and stratified according to the genetic background of the HepG2 cells

| | **ER width** | **ANOVA** | **Student t-test** | ***P-value**** |
|---|---|---|---|---|
| Cas9$^{+}$ | 0.09 {0.07-0.13} | <0.0001 | | |
| MBOAT7$^{-/-}$ | 0.18 {0.14-0.25} | <0.0001 | <0.0001 | **<0.0001** |
| TM6SF2$^{-/-}$ | 0.30 {0.23-0.37} | <0.0001 | <0.0001 | **<0.0001** |
| MBOAT7$^{-/-}$TM6SF2$^{-/-}$ | 0.35 {0.27-0.44} | <0.0001 | <0.0001 | **<0.0001** |

Values are reported as median {IQR}. **P*-values are adjusted at *post hoc* Dunn's multiple comparison test and compared to Cas9$^{+}$ control group.

**Table S5**. Sequence of primers used in quantitative real-time PCR experiments.

| | **Forward 5'→3'** | **Reverse 5'→3'** |
|---|---|---|
| *ATF4* | AAACCTCATGGGTTCTCCAG | GGCATGGTTTCCAGGTCACT |
| *ATF6* | AATTCTCAGCTGATGGCTGT | TGGAGGATCCTGGTGTCCAT |
| *GRP78* | CTTGCCGTTCAAGGTGGTTG | CTGCCGTAGGCTCGTTGAT |
| *MnSOD2* | CAAATTGCTGCTTGTCCAAA | TCTTGCTGGGATCATTAGGG |
| *PPARα* | ATGGCATCCAGAACAAGGAG | TCCCGTCTTTGTTCATCACA |
| *MnSOD2* | CAAATTGCTGCTTGTCCAAA | TCTTGCTGGGATCATTAGGG |
| **TM6SF2 exon I* | GGCTGCCTATGCTCTCACCTT | TGCCTCCAGCAAACACCAA |
| **U6-sgRNA* | TTTCTTGGGTAGTTTGCAGTTTT | CGACTCGGTGCCACTTTT |
| *XBP1* | GAAGCCAAGGGGAATGAAGT | GCCCAACAGGATATCAGACTC |
| *β-actin* | GCTACAGCTTCACCACCACA | AAGGAAGGCTGGAAAAGAGC |

*human primers used for Sanger sequencing

**Table S6**. List of TaqMan Probes used in quantitative real-time PCR experiments.

| Probes | Catalog Number |
|---|---|
| TM6SF2 | ThermoFisher #Hs00403495_m1 |
| MBOAT7 | ThermoFisher #Hs00383302_m1 |
| GCK | ThermoFisher #Hs01564555_m1 |
| PFK-L | ThermoFisher #Hs01036347_m1 |
| G3PDH | ThermoFisher #Hs02786621_g1 |

**Table S7**. List of antibodies and relative dilutions used in Western blotting and ICC experiments.

| **Antibody** | **Catalog Number** |
|---|---|
| TM6SF2 (1:1000 WB) | ThermoFisher # PA5-69304 |
| MBOAT7 (1:500 WB) | Sigma-Aldrich #AV49811 |
| ApoB-100 | ThermoFisher #HYB0690202 |
| P(Ser473) -Akt (1:1000 WB) | Cell signaling #9271S |
| Akt (1:1000 WB) | Cell signaling #2938S |
| P(Thr37/46) -4E-BP1 (1:1000 WB) | Cell signaling #2855S |
| 4E-BP1 (1:1000 WB) | Cell signaling #9644S |
| P(Ser2448) -mTOR (1:1000 WB) | Cell signaling #5536S |
| mTOR (1:1000 WB) | Cell signaling #2983S |
| PGC1-α | Novus #NBP1-04676 |
| COX-I | Abcam #ab110216 |
| SDHA | Abcam #ab110216 |
| β-actin (1:5000 WB) | Abcam #ab6276 |

#### Supplemental figure legends

**Figure S1**: *The co-presence of the PNPLA3 rs738409, TM6SF2 rs58542926 and MBOAT7 rs641738 variants predicted the risk to develop NAFLD and hepatic fibrosis.* **A**) Multivariate ordinal regression analysis adjusted for age, gender, BMI and T2D, the co-presence of I148M PNPLA3, E167K TM6SF2 and the rs641738 in *MBOAT7* variants associated with NAFLD activity score (NAS). At nominal logistic regression analysis adjusted for age, gender, BMI and T2D, the co-presence of 3 SNPs increased the risk to develop NAFLD (**B**), mild fibrosis (**C**) and advanced fibrosis (**D**). *0: indicates the absence of risk variants; 1-2-3 indicates the total number of risk variants carried.

**Figure S2:** *CRISPR/Cas9-mediated TM6SF2 ablation in HepG2 cells.* **A**) A schematic representation of the *TM6SF2* sequence (Gene ID: 53345; referred to as Transcript variant 1: NM_001001524.3) highlighted the same clonal Cas9-induced indel mutations in both TM6SF2$^{-/-}$ and MBOAT7$^{-/-}$TM6SF2$^{-/-}$ clones (blue) of 202 nucleotides (Δ202). Cas9 cutting site is indicated by the symbol '|' (red) and the transcription start site (ATG) of the protein coding sequence (NP_001001524.2) is in green. **B**) mRNA and protein expression of TM6SF2 was evaluated through qRT-PCR and Western blot, respectively. TM6SF2 reduction was detected in TM6SF2$^{-/-}$ and MBOAT7$^{-/-}$ TM6SF2$^{-/-}$ cells (adjusted* p<0.05 and **p<0.01 *vs* Cas9$^{+}$ and *vs* MBOAT7$^{-/-}$). **C**) MBOAT7 mRNA and protein levels were lower in MBOAT7$^{-/-}$ and MBOAT7$^{-/-}$ TM6SF2$^{-/-}$cells compared to Cas9$^{+}$ and TM6SF2$^{-/-}$cells (adjusted *p<0.05and **p<0.01, respectively). **D**) Apolipoprotein B-100 (ApoB) protein was assessed in cell supernatants by Western blot and normalized to the entire lane of the Ponceau stain. Either TM6SF2$^{-/-}$ and MBOAT7$^{-/-}$ TM6SF2$^{-/-}$ showed low ApoB levels (*p<0.05 *vs* Cas9$^{+}$ and **p<0.01 *vs* MBOAT7$^{-/-}$ at Two-tailed Student *t*-test). **E**) TAG-rich lipoproteins' secretion has been measured in cell supernatants and normalized to levels of total cholesterol by using Cholesterol Colorimetric Assay Kit - HDL and LDL/VLDL (Abcam, Cambridge, UK). Both TM6SF2$^{-/-}$ and MBOAT7$^{-/-}$ TM6SF2$^{-/-}$ dampened TAG-rich lipoproteins' release (adjusted *p<0.05 *vs* control and **p<0.01 *vs* MBOAT7$^{-/-}$). Data was normalized to β-actin housekeeping gene for qRT-PCR and Western blot and expressed as fold increase (Arbitrary Units - AU). At least three independent experiments were conducted.

**Figure S3**: *TM6SF2 deletion induced spontaneous LDs accumulation.* **A**) Spontaneous development of LDs in TM6SF2$^{-/-}$ and MBOAT7$^{-/-}$TM6SF2$^{-/-}$ cells was assessed by ORO staining (200X magnification). **B)** Alkaline phosphatase (ALP) stained on LDs surface in MBOAT7$^{-/-}$, TM6SF2$^{-/-}$ and MBOAT7$^{-/-}$TM6SF2$^{-/-}$ clones, highlighting in yellow differences in LDs size for each condition. **C**) Representative TEM images obtained by ultrathin 70 nm sections of hepatocytes. Micrographs highlighted LDs accumulation and dimensions in MBOAT7$^{-/-}$, TM6SF2$^{-/-}$ and MBOAT7$^{-/-}$TM6SF2$^{-/-}$ cells and were acquired at different bar scale length (bar scale: 1 µm). Black arrows indicated LDs, whereas organelles are referred to as capital letters (N: nucleus; ER: endoplasmic reticulum; M: mitochondria). **D**) Heatmap of TAG species was generated by calculating log2 fold change (log2FC) ratio between MBOAT7$^{-/-}$/Cas9$^{+}$ quantification. Red and blue boxes indicate overexpression or repression, respectively. **E**) Principal-component analysis (PCA) of lipidomic profile of MBOAT7$^{-/-}$, TM6SF2$^{-/-}$ and MBOAT7$^{-/-}$TM6SF2$^{-/-}$ models. **F-G**) Relatively enriched DAGs and

TAGs in $MBOAT7^{-/-}$ $TM6SF2^{-/-}$ cells compared to $MBOAT7^{-/-}$. Data are expressed as percentage mean (%) of DAG or TAG species and standard deviation (SD). adjusted *p<0.05 or **p<0.01 *vs* $MBOAT7^{-/-}$.

**Figure S4:** *The impact lyso-phosphatidylcholines (Lyso-PCs) metabolism in TM6SF2-silenced cells.* **A**) Heatmap of PCs was generated by calculating log2 fold change (log2FC) ratio between $MBOAT7^{-/-}$/$Cas9^{+}$ quantification. Red and blue boxes indicate overexpression or repression, respectively. **B-C**) Relative enrichment of Lyso-PCs in $TM6SF2^{-/-}$ and $MBOAT7^{-/-}$ $TM6SF2^{-/-}$ cells *vs* $Cas9^{+}$. **D-E**) Relative enrichment of Lyso-PC and PC species in $MBOAT7^{-/-}$ $TM6SF2^{-/-}$ cells *vs* $MBOAT7^{-/-}$. Data are expressed as percentage mean (%) of Lyso-PCs and standard deviation (SD). adjusted *p<0.05 or **p<0.01 *vs* $Cas9^{+}$ and *vs* $MBOAT7^{-/-}$.

**Figure S5:** *TM6SF2 silencing induced mitochondrial oxidative damage.* **A**) The $H_2O_2$ levels were measured in cell lysates (adjusted *p<0.05 and **p<0.01 vs $Cas9^{+}$) through DCF ROS/RNS Colorimetric Assay Kit (Abcam, Cambridge, UK). **B**) The MnSOD2 mRNA expression was assessed by qRT-PCR and normalized to β-actin housekeeping gene (adjusted **p<0.01 *vs* $Cas9^{+}$ and *vs* $MBOAT7^{-/-}$). **C**) The malondialdehyde (MDA) secretion was colorimetrically measured in cell supernatants following the manufacturer's instruction (adjusted *p<0.05 *vs* $Cas9^{+}$). **D**) PPARα mRNA expression was assessed by qRT-PCR and normalized to β-actin housekeeping gene (adjusted *p<0.05 *vs* $MBOAT7^{-/-}$ and *vs* $TM6SF2^{-/-}$; adjusted **p<0.01 *vs* $Cas9^{+}$). Data are expressed as fold increase (Arbitrary Unit-AU) compared to control group. At least three independent experiments were conducted. adjusted *p<0.05 or **p<0.01 *vs* $Cas9^{+}$ and *vs* $MBOAT7^{-/-}$.

**Figure S6:** *$TM6SF2^{-/-}$ and $MBOAT7^{-/-}TM6SF2^{-/-}$ cells aberrantly activated PI3K/Akt/mTOR pathway.* **A**) Phosphorylation of Akt at serine 437 residue (P(S437)-Akt), mTOR at serine 2448 residue (p(S2448)-mTOR), phosphorylation of 4E-BP1 at threonine 37/46 residues P(Thr37/46)-4E-BP1 and total Akt, mTOR and 4E-BP1 were evaluated by Western blot. **B-D)** Quantification of P(S437)-Akt/total Akt, p(S2448)-mTOR/total mTOR, P(Thr37/46)-4E-BP1/total 4E-BP1 ratios were measured through ImageJ software and normalized to β-actin housekeeping gene. **E**) Cells were exposed to Sorafenib (5 µM) and cell growth was monitored through MTS assay for 0-24-48-72 hrs and 1 week. $MBOAT7^{-/-}$ $TM6SF2^{-/-}$ cells showed a significant resistance to Sorafenib cytotoxicity at 24 and 48 hrs (adjusted **p<0.01 vs $Cas9^{+}$). MTS absorbance (λ=490 nm) was recorded at 0-24-48-72 hrs and 1 week. **F**) Representative images of wound healing assay were acquired at 0-24-48 hrs (100X magnification) upon Sorafenib (5 µM) administration. The dotted lines indicate the scratch width. Data are expressed as fold increase (Arbitrary Unit-AU) compared to control group. At least three independent experiments were conducted. adjusted *p<0.05 or **p<0.01 *vs* $Cas9^{+}$ and $MBOAT7^{-/-}$.
